# Rad24-RFC loads the 9-1-1 clamp by inserting DNA from the top of a wide-open ring, opposite the mechanism of RFC/PCNA

**DOI:** 10.1101/2021.10.01.462756

**Authors:** Fengwei Zheng, Roxana E. Georgescu, Nina Y. Yao, Michael E. O’Donnell, Huilin Li

**Affiliations:** Department of Structural Biology, Van Andel Institute, Grand Rapids, Michigan, USA; DNA Replication Laboratory, The Rockefeller University, New York, New York, USA; Howard Hughes Medical Institute, The Rockefeller University, New York, New York, USA

## Abstract

In response to DNA damage, the ring-shaped 9-1-1 clamp is loaded onto 5’ recessed DNA to arrest the cell cycle and activate the DNA damage checkpoint. The 9-1-1 clamp is a heterotrimeric ring that is loaded in *S. cerevisiae* by Rad24-RFC, an alternative clamp loader in which Rad24 replaces the Rfc1 subunit in the RFC1-5 clamp loader of PCNA. Unlike RFC that loads the PCNA ring onto a 3’-ss/ds DNA junction, Rad24-RFC loads the 9-1-1 ring onto a 5’-ss/ds DNA junction, a consequence of DNA damage. The underlying 9-1-1 clamp loading mechanism has been a mystery. Here we report two 3.2-Å cryo-EM structures of Rad24-RFC bound to DNA and either a closed or 27 Å open 9-1-1 clamp. The structures reveal a completely unexpected mechanism by which a clamp can be loaded onto DNA. The Rad24 subunit specifically recognizes the 5’-DNA junction and holds ds DNA outside the clamp loader and above the plane of the 9-1-1 ring, rather than holding DNA inside and below the clamp as in RFC. The 3’ ssDNA overhang is required to obtain the structure, and thus confers a second DNA binding site. The bipartite DNA binding by Rad24-RFC suggests that ssDNA may be flipped into the open 9-1-1 ring, similar to ORC-Cdc6 that loads the Mcm2-7 ring on DNA. We propose that entry of ssDNA through the 9-1-1 ring triggers the ATP hydrolysis and release of the Rad24-RFC. The key DNA binding residues are conserved in higher eukaryotes, and thus the 9-1-1 clamp loading mechanism likely generalizes.

## INTRODUCTION

The DNA damage response is essential for maintaining genome integrity in all eukaryotes (Harrison and Haber, 2006; Sancar et al., 2004; Su, 2006; Zhou and Elledge, 2000). In response to DNA damage or replication stress, a checkpoint signaling pathway is activated to arrest the cell cycle and initiate repair of the DNA damage, or trigger apoptosis (Abraham, 2001; Donehower, 2014; Kumagai et al., 1998; Patil et al., 2013; Peng et al., 1997; Walworth et al., 1993). Upon DNA damage, single-strand (ss) DNA is produced by CMG helicase advance that continues after replicative DNA polymerases stall at sites of DNA damage. These ssDNA gaps contain 5’ ss/double-strand (ds) DNA junctions (referred to here as a “5’-DNA junction”), which are normally very few and transient during replication. The 5’-DNA junctions provide “targets of opportunity” to signal that DNA damage has occurred. In both yeast and human, the DNA damage response is initiated by the loading of the checkpoint complex, Rad9–Hus1–Rad1 in human or Ddc1–Mec3–Rad17 in yeast (both referred to as 9-1-1; **Fig. 1a**), onto 5’-DNA junctions produced as a consequence of DNA damage (Ellison and Stillman, 2003; Majka et al., 2006; Majka and Burgers, 2003). The 9-1-1 clamp loading is accomplished by a specialized clamp loader called RAD17-RFC (Replication Factor C) in human and Rad24-RFC in *S. cerevisiae* (Parrilla-Castellar et al., 2004).

**Fig. 1.**
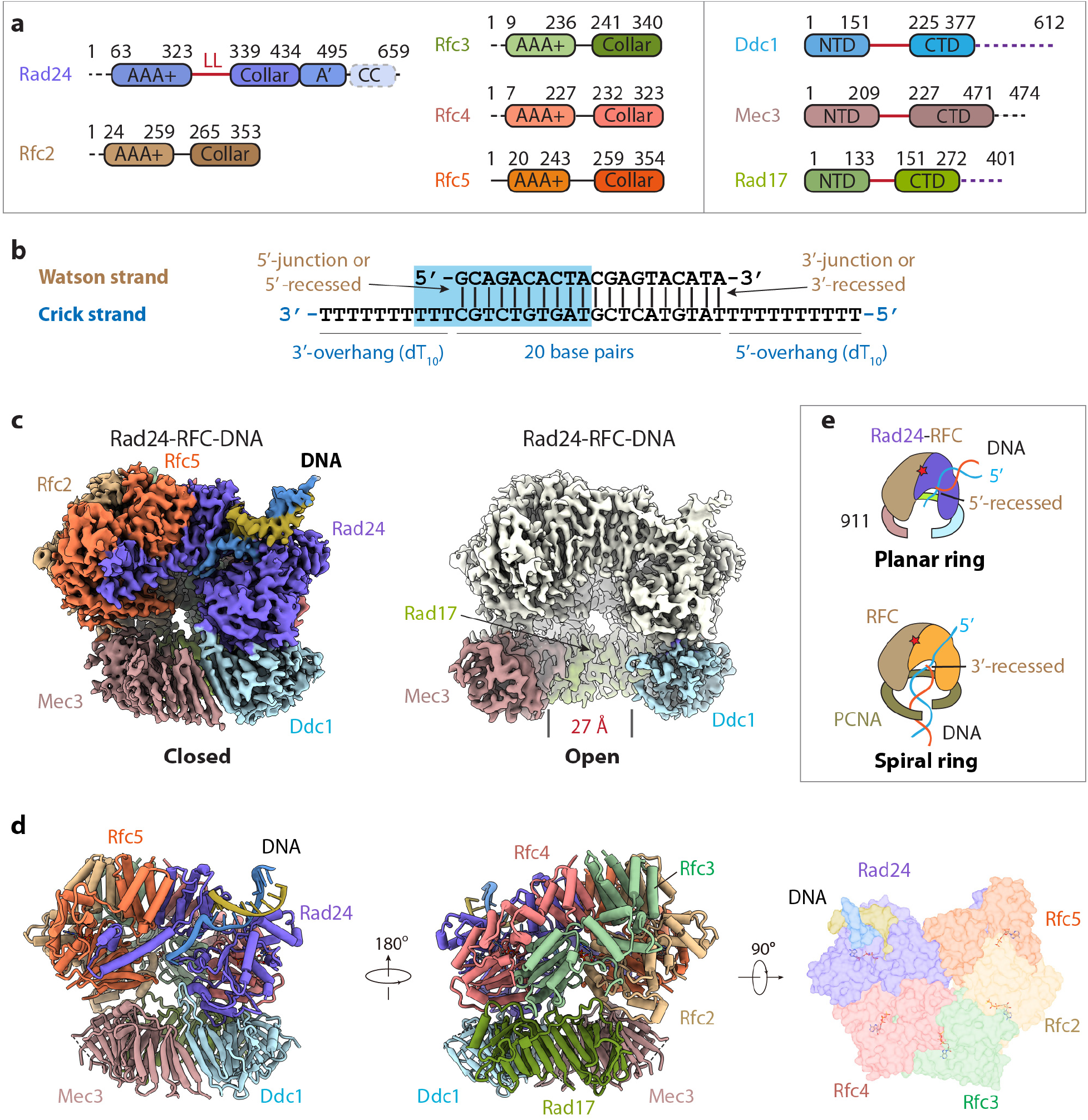
Cryo-EM 3D maps of the S.c. Rad24-RFC–9-1-1-DNA ternary complex. **a**) Domain architecture of the eight proteins of the Rad24-RFC–9-1-1 complex: the clamp loader proteins Rad24 and Rfc2-5, and the 9-1-1 clamp proteins Ddc1, Mec3 and Rad17. The Rad24 A’ domain is associated with the collar domain that is also present in Rfc1 which Rad24 replaces. The unique Rad24 C-terminal coiled coil (CC) is disordered. The dashed lines indicate unsolved regions. LL is the long linker between the Rad24 AAA+ and collar domains. The red lines between NTDs and CTDs of 9-1-1 subunits represent the inter-domain connecting loop (IDCL), and the dashed purple lines represent the long C-terminal tails of Ddc1 and Rad17. **b**) DNA substrate used to form the ternary complex. Both the 5’-DNA junction and 3’-DNA junction are present. Nucleotides in the cyan box are resolved in the structure. **c**) Segmented 3D maps of the Rad24-RFC–9-1-1 clamp–DNA complex in the closed (left) and open (right) states. Maps are surface rendered at 0.1 threshold, except for Ddc1 in the open conformation, which is separately displayed at 0.07 threshold. **d**) Model of the closed state complex in the front and back view (cartoon) and in the top view (surface). **e**) A sketch comparing the DNA binding mode of the Rad24-RFC–9-1-1 (upper panel, this study) with that of the RFC–PCNA, based on the T4 clamp–clamp loader–DNA structure (lower panel; PDB code 3U60). Note the drastically different DNA positions and the spiral versus planar rings of the two systems.

The 9-1-1 complex is a PCNA (proliferating cell nuclear antigen)-like ring conserved from yeast to human (Hang et al., 1998; Lieberman et al., 1992; Lieberman et al., 1996; Venclovas and Thelen, 2000), and is able to slide on DNA (Ellison and Stillman, 2003; Majka and Burgers, 2003). Three crystal structures published in 2009 showed that the human 9-1-1 clamp is a heterotrimeric ring-shaped DNA clamp (Dore et al., 2009; Sohn and Cho, 2009; Xu et al., 2009). Each 9-1-1 subunit has a PCNA-like core consisting of an N-terminal domain (NTD) and a C-terminal domain (CTD) that are connected by a long inter-domain connecting loop (IDCL). The surface of the inner chamber of the 9-1-1 ring is lined by basic residues that facilitate DNA binding and/or sliding. The human 9-1-1 subunit Rad9 (Ddc1 in yeast) has a very long and flexible C-tail with several phosphorylation sites. The phosphorylated Rad9/Ddc1 C-tail recruits TopBP1 to activate the checkpoint kinase ATR (ataxia telangiectasia mutated, ATM, and Rad-3 related; Mec1 in yeast) (Delacroix et al., 2007; Lee et al., 2007). The active kinase goes on to phosphorylate and activate the checkpoint kinase 1 (Enders, 2008; Tannous et al., 2021). This further activates the downstream checkpoint proteins, leading to the arrest of the cell cycle progression (Abraham, 2001; Zhou and Elledge, 2000).

Despite their similar ring structure and the shared ability to serve as a platform to coordinate numerous proteins involved in DNA metabolism, 9-1-1 and PCNA function differently. PCNA is a homotrimer that functions as a sliding clamp for the high-fidelity replicative DNA polymerases to achieve high processivity, and also interacts with numerous other types of DNA polymerases and a wide variety of DNA metabolic factors (Bruck and O’Donnell, 2001; Kelman, 1997; Lancey et al., 2020; Madru et al., 2020; Zheng et al., 2020). The PCNA clamp is loaded onto the 3’-recessed template/primer DNA junction (i.e., “3’-DNA junction”) by the RFC loader (Kelch et al., 2012). In contrast, the alternative Rad17(Rad24)-RFC clamp loader of 9-1-1 is specific for the 5’-DNA junction. Rad17(Rad24)-RFC is one of three alternative RFC-like complexes in eukaryotes in which Rfc1 is replaced by another subunit (Kim and MacNeill, 2003). Both RFC and Rad17(Rad24)-RFC clamp loaders are specialized heteropentameric AAA+ (ATPases Associated with diverse cellular Activities) complexes. RFC is composed of Rfc subunits 1-5 and loads PCNA onto DNA. Specific to this report, *S. cerevisiae* Rad24-RFC is formed by the replacement of the largest subunit of RFC (Rfc1) with Rad24 (RAD17 in human) and loads the DNA damage 9-1-1 ring onto DNA to signal DNA damage (**Fig. 1a**).

The Rad17(Rad24)-RFC has been demonstrated to load the 9-1-1 clamp onto a 5’-junction in vitro (Ellison and Stillman, 2003; Majka et al., 2006). The 9-1-1 clamp loading process is assisted by RPA (Replication Protein A) that covers the ssDNA adjacent to the 5’-DNA junction (Zou et al., 2003). In this connection, the Rad24-RFC is reported to require RPA bound to the exposed ssDNA for directional loading onto a 5’ versus a 3’ DNA junction (Majka et al., 2006). RPA was also shown to interact with the C-terminal coiled coil of Rad24 to facilitate 9-1-1 loading by Rad24-RFC in a 5’-junction specific manner (Piya et al., 2015). Furthermore, earlier studies document that the large subunit of RPA (Rpa1) binds directly to Rfc4 and Rfc5 which are present in the Rad24-RFC loader (Kim and Brill, 2001; Naiki et al., 2000; Shimomura et al., 1998; Yuzhakov et al., 1999).

The structures of the yeast and human RFC–PCNA complexes in the absence of DNA showed that each Rfc subunit contains the AAA+ two domain module (referred to here as the AAA+ domain) and a C-terminal collar domain (Bowman et al., 2004; Gaubitz et al., 2020; Indiani and O’Donnell, 2006). The RFC pentamer is configured in a right-hand spiral consisting of two tiers, with five collar domains forming a sealed ring at the top tier, and five AAA+ regions forming a spiral with a gap between Rfc1 and Rfc5 at the lower tier (Jeruzalmi et al., 2002; Kelch, 2016; O’Donnell and Kuriyan, 2006). Rfc1 has an additional A’ domain associated with the collar domain that connects with the adjacent Rfc5 AAA+ domain, thereby partially filling the gap between Rfc1 and Rfc5. The A’ domain is also present in Rad24. The ATP binding sites of the RFC loaders are located at the interfaces of two adjacent AAA+ domains, with one AAA+ domain binding the ATP via the conserved Walker A (P-loop) and Walker B (DExx box) motifs, and the neighboring AAA+ domain contributing an arginine finger within a conserved serine-arginine-cysteine (SRC) motif, and a central helix motif (Jeruzalmi et al., 2002; Kelch, 2016; Kelch et al., 2012; O’Donnell and Kuriyan, 2006). ATP hydrolysis in these clamp loaders occurs in *trans,* stimulated by the arginine finger of the adjacent subunit. However, the Rfc1 subunit – as well as Rad24 – does not have the SRC motif, nor do they have the central helix motif that are conserved in Rfc2-5; features that are also conserved in the bacterial clamp loader γ_3_δδ’ complex (Jeruzalmi et al., 2001; Kelch et al., 2011). So far, the high-resolution structure of a DNA-bound RFC-PCNA has not been achieved (Miyata et al., 2005), and the structure of a DNA-bound clamp-clamp loader complex is available only for the T4 bacterial phage system in which both the clamp loader and the clamp assume a spiral shape matching the helical symmetry of the DNA (Kelch et al., 2011).

There have been a few low-resolution EM observations of the human Rad17–RFC-9-1-1 and the yeast Rad24-RFC–9-1-1 complexes, revealing the ring-like shapes of the 9-1-1 clamp and the loaders (Bermudez et al., 2003; Griffith et al., 2002; Liu, 2019; Shiomi et al., 2002). Due to the lack of a high-resolution structure, it has been unclear if the 9-1-1 clamp is indeed loaded onto the 5’-DNA junction, and if true, how could Rad24-RFC load the 9-1-1 clamp ring in an essentially opposing direction of the RFC loading of the PCNA ring? Here we describe the first Rad24-RFC–9-1-1 clamp structures bound to DNA and nucleotides, in both closed and open conformation, at a resolution of 3.2 Å. Our structures revealed a surprising DNA binding mode, explaining the unique 9-1-1 clamp loading mechanism that is drastically different from that of PCNA loading by RFC.

## RESULTS AND DISCUSSION

### 1. In vitro assembly and cryo-EM of the Rad24-RFC–9-1-1–DNA ternary complex

We separately expressed and purified the Rad24-RFC from the Baker’s yeast and the 9-1-1 clamp from *E coli* (**Extended Data Fig. 1a**). We first examined in vitro binding of a 3’-recessed DNA substrate and found by cryo-EM that this DNA failed to induce the assembly of the Rad24–RFC-9-1-1–DNA ternary complex (**Extended Data Fig. 1b-c**). Next, we designed a two-tailed DNA substrate in which both a 3’-junction and a 5’-junction are present (**Fig. 1b**). This DNA substrate enables us to determine in an unbiased manner which recessed end Rad24-RFC prefers to bind in the absence of RPA. We mixed the purified proteins with the DNA substrate in the presence of 0.5 mM ATPgS, a weakly hydrolyzable ATP analog. Cryo-EM analysis indicated a successful assembly of the Rad24-RFC–9-1-1–DNA ternary complex (**Extended Data Fig. 1d-e**). Subsequent 2D and 3D classification, 3D reconstruction, and 3D variability analysis led to the capture of two conformations of the ternary complex at 3.2 Å average resolution (**Extended Data Figs. 2-4, Extended Data Table 1**). In one conformation the 9-1-1 ring is closed while in the other the ring is wide open with a 27-Å gap (**Fig. 1c, Supplementary Video 1**). The Rad24 structure was built *de novo* while modeling of Rfc2-5 referenced the shared subunits in the RFC-PCNA crystal structure (Bowman et al., 2004). The EM density of the 9-1-1 ring had a slightly lower resolution of about 3.8 Å with Mec3 having the best density (**Extended Data Fig. 3**). The yeast 9-1-1 subunits contain long and disordered insertion loops and are larger than the yeast PCNA (276 residues) or the human 9-1-1 proteins (Venclovas and Thelen, 2000) (**Fig. 1a**), but they all have a core structure conserved with the PCNA and human 9-1-1 clamps, facilitating modeling of the yeast 9-1-1 core structure (Bowman et al., 2004; Gaubitz et al., 2020). The EM density of the DNA substrate was of sufficiently high resolution to distinguish purine from pyrimidine, enabling atomic modeling and unambiguous sequence assignment (**Extended Data Fig. 4**).

In the atomic model of the Rad24-RFC–9-1-1–DNA complex, the five clamp loader subunits are arranged counterclockwise Rad24-Rfc4-Rfc3-Rfc2-Rfc5 when viewed from the top (**Fig. 1d**), and importantly, only the 5’-recessed end-containing half of the DNA was visible and stably bound to Rad24-RFC (i.e. the 3’-recessed end-containing half of the DNA was not visible). The resolved DNA includes 10-bp of the 5’-DNA junction and three ssDNA nucleotides of the 3’-overhang (**Fig. 1b, d**). The DNA was cradled between the AAA+ domain and collar domain of Rad24 in a manner that is opposite of DNA binding in the RFC-PCNA complex (Simonetta et al., 2009) (**Fig. 1d, e**). Because our DNA substrate contains both 3’- and 5’-recessed ends, the fact that Rad24-RFC bound only to the 5’-recessed half of the substrate confirms the reported preference of the Rad24-RFC (Rad17-RFC in human) to load 9-1-1 onto a 5’-recessed DNA end (Ellison and Stillman, 2003; Majka et al., 2006), and appears to do so even in the absence of RPA under the conditions used here.

### 2. Unique features of Rad24 ensure specific recognition of the 5’-DNA junction

Here we will use the closed state to illustrate the DNA binding, although the DNA binding mode is similar in both open and closed states (**Supplementary Video 2**). The most striking feature is that the DNA is bound only by the Rad24 subunit in an extended groove between its AAA+ and collar domains (**Fig. 2a**). Several features of Rad24 apparently have enabled the unique DNA binding mode in the Rad24-RFC. First is the 15-residue “long linker” (sometimes referred here as “LL”), from residue 324 to 338 that connects the Rad24 AAA+ module to the collar domain (**Figs. 1a and 2a**). The long linker has enabled the lid and Rossmann fold domains (in the AAA+ module) to move 35 Å and rotate 20° away from the Rad24 collar domain, forming the DNA-binding cleft (**Supplementary Video 3**). This linker is one-residue long in Rfc1, the Rad24 equivalent of the RFC complex (Bowman et al., 2004), and is 21-residues long (residue 827 to 847) but disordered in the human RFC1 (Gaubitz et al., 2020) (**Extended Data Fig. 5**). In both yeast Rfc1 and human RFC1, the collar domain is next to the lid and Rossmann fold domains such that no DNA binding groove exists (**Extended Data Fig. 6**).

**Fig. 2.**
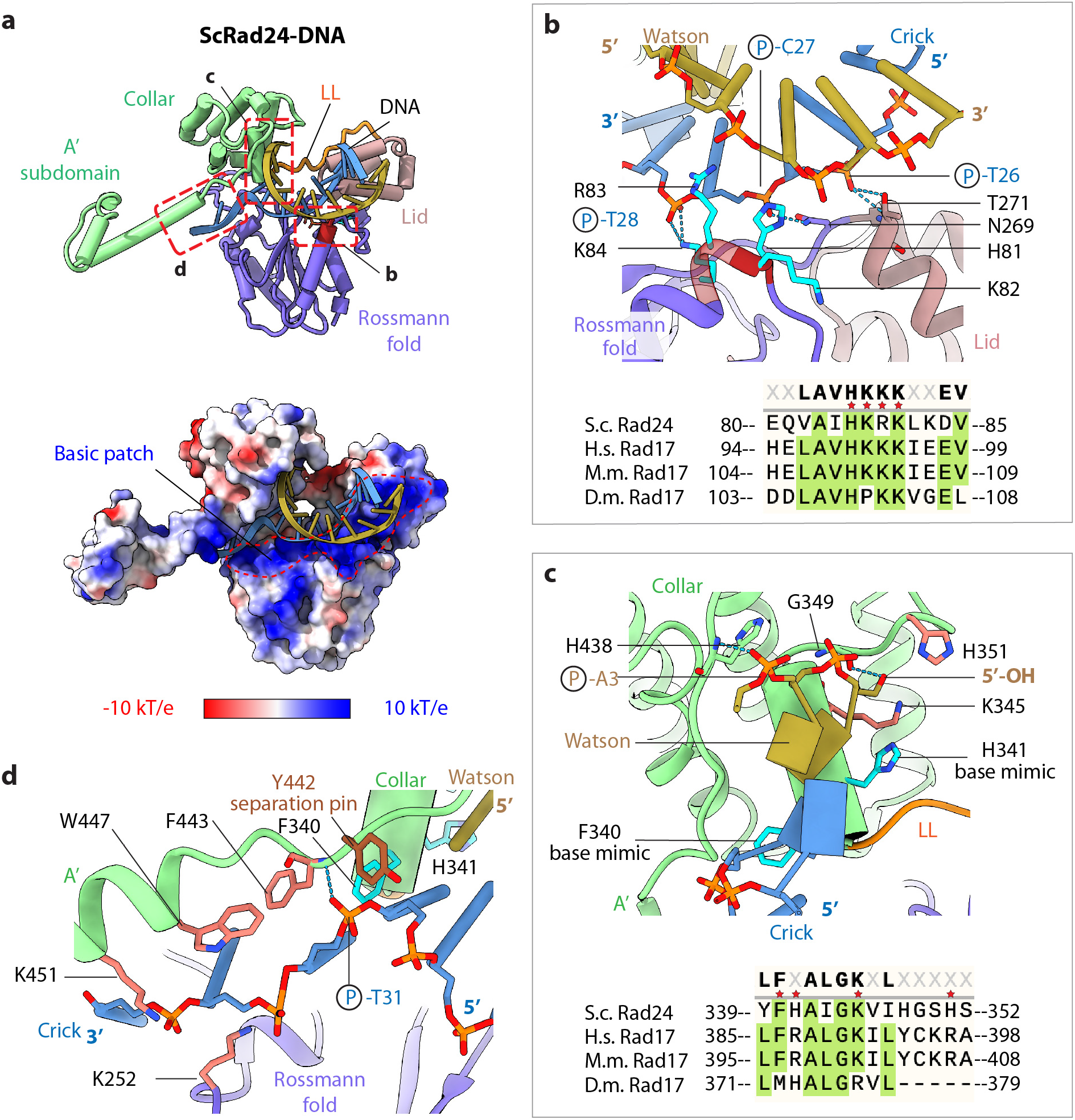
Rad24 interaction with the 5’-DNA junction. **a**) Top: Structure of Rad24 bound to the DNA in cartoon view. The Watson strand is orange, and the Crick strand is cyan. The four basic residues (81-HKRK-84) in the AAA+ domain are painted dark red. The areas marked by three dashed red rectangles are close-up views in panels b, c, and d, respectively. Bottom: Electrostatic surface view showing a contiguous basic patch on the top surface of the AAA+ domain that binds DNA. LL: Long linker between AAA+ and collar domains that enables the formation of a large groove to accommodate the DNA. **b**) Top: Interaction between the AAA+ domain and DNA. The four tandem basic residues are shown in cyan sticks. His-81 and Arg-83 insert into the DNA minor groove, and Lys-84 forms two H-bonds with the DNA T28 phosphate of the Crick strand. Asn-269 of AAA+ domain and Thr-271 of the lid domain form three H-bonds with the DNA phosphates of C27 and T26. Bottom: Sequence alignment of the Rad24 four tandem basic residues. S.c., *Saccharomyces cerevisiae;* H.s., *Homo sapiens;* M.m., *Mus musculus;* and D.m., *Drosophila melanogaster.* **c)** Top: The Rad24 collar domain stabilizes the 5’-DNA junction. Phe-340 and His-341 mimic a nucleotide base and form a hydrophobic stack with the last base pair at the 5’-junction; they each rotate 32° with respect to the interacting base, resembling the 36° rotation of a normal base. The basic His341, Lys-345, and His-351 surround the 5’-OH of the Watson strand and prevent a 5’-overhang ssDNA from binding there. 5’-OH and the A3 phosphate form H-bonds with Gly-349 and His-438, respectively. Bottom: Sequence alignment of Rad24 base mimicking residue F340 and the 5’-OH blocking H341 residue. **d)** Residues guiding ssDNA 3’-overhang into the Rad24-RFC chamber. Tyr-442 is positioned at the 5’-DNA junction resembling a DNA separation pin.

The second important feature of Rad24 is the presence of a long and contiguous basic path on the top of the AAA+ domain, enabling stable DNA binding (**Fig. 2a**). Such a continuous basic patch is absent in the yeast Rfc1 and human RFC1 (**Extended Data Fig. 6**). The basic path is comprised of four tandem positively charged residues (81-HKRK-84) in the AAA+ domain (**Fig. 2b**). The side chains of His-81 and Arg-83 insert into the DNA minor groove, and Lys-84 forms two hydrogen bonds with the T-28 phosphate of the Crick strand. Lys-82 points away from DNA but contributes to the overall positively charged environment of the DNA path. In addition, Asn-269 of the AAA+ domain and Thr-271 of the Lid domain contact the DNA, forming hydrogen bonds with the phosphate groups of C-27 and T-26 of the Crick strand. The four-residue basic patch is largely conserved among the 9-1-1 loading eukaryotic Rad24/Rad17-RFCs but is not present in the PCNA loading Rfc1-RFC (**Fig. 2d**, **Extended Data Fig. 5**).

At the 5’-junction, the aromatic Phe-340 and His-341 in the Rad24 α-helix 11 form hydrophobic stacking interactions with the DNA residues C-30 and G-1 (see the secondary structure assignment in **Extended Data Fig. 5**), respectively, thereby stabilizing the last base pair G-1:C-30 (**Fig. 2c**). These two aromatic side chains are rotated ~32° relative to the last two bases (G-1 and C-30), mimicking the normal 36° rotation of one bp to another bp in dsDNA. In vitro synthesized DNA are generally capped at the 5’ end by a hydroxyl group (OH) rather than a phosphate, and indeed, DNA substrates with a 5’-OH have been used for in vitro 9-1-1 clamp DNA binding and/or loading assays (Ellison and Stillman, 2003; Majka et al., 2006). Importantly, the G1 5’-OH of the in vitro synthesized DNA in our structure is surrounded by three positively charged residues His-341, Lys-345, and His-351. These residues are well positioned to neutralize the phosphate at the 5’-DNA junction and thus prevent a 5’-overhang from binding at the site. The 5’-junction is further stabilized by a H-bond between Gly-349 of the collar domain and 5’-OH of DNA and by another H-bond between His-438 of the A’ subdomain of Rad24 and the A3 phosphate of the DNA (**Fig. 2c**). These structural features likely account for Rad24’s specificity for the 5’-DNA junction (Ellison and Stillman, 2003; Majka et al., 2006).

Beyond the 5’-DNA junction, Rad24 further stabilizes the first three nucleotides (T31, T32, and T33) of the 3’-overhang (**Fig. 2e**). Specifically, Tyr-442, structurally equivalent to the “separation pin” Tyr-316 in the *E. coli* clamp loader (Simonetta et al., 2009), is positioned right at the 5’-junction of the ssDNA and dsDNA. Then, Phe-443, Trp-447, and Lys-451 from the Rad24 A’ subdomain stabilize the exposed bases of the three 3’-overhang nucleotides. Finally, Lys-252 forms electrostatic interaction with the DNA T32 phosphate, and the mainchain nitrogen of Phe-443 forms a H-bond with the DNA T31 phosphate of the Crick strand DNA, guiding the 3’ ssDNA overhang towards the interior chamber of the Rad24-RFC loader. These DNA-interacting residues are conserved among yeast Rad24 and metazoan Rad17 (**Fig. 2f**).

### 3. Rad24-RFC is poised to hydrolyze ATP

Each of the five nucleotide-binding sites of the Rad24-RFC loader is occupied by a nucleotide. In both closed and open 911 clamp states, the first four interfaces (Rad24:Rfc4, Rfc4:Rfc3, Rfc3:Rfc2, and Rfc2:fc5) are occupied by ATPγS and Mg^2+^, while an ADP occupied the last interface between Rfc5 and the A’ subdomain of Rad24 (**Fig. 3a**). This observation suggests that the 9-1-1 gate opening does not require ATP hydrolysis by Rad24-RFC. It is remarkable that the nucleotide binding pattern resembles those in the RFC–PCNA structures in which the first four interfaces bound to ATPγS and the last interface bound to ADP (Bowman et al., 2004; Gaubitz et al., 2020), and this is also the case in the T4 clamp-clamp loader-DNA structure (Kelch et al., 2011). Sequence alignment identified conserved ATPase motifs such as the Walker A (P-loop; consensus: GxxGxGK[T/S], where x is any residue), Walker B (DExx box), Sensor 1, and Sensor 2 motifs (Jeruzalmi et al., 2002; Kelch et al., 2012) (**Extended Data Figs. 5, 7).** Starting from Rad24 and counterclockwise, in the first four nucleotide binding pockets, the Walker A motif (P-loop, see sequence alignment) wraps the triphosphate, and the P-loop lysine forms an H-bond with the phosphate group; the first acidic D/E of the Walker B motif (DExx box) coordinates a Mg^2+^ ion and H-bonds with ATPγS, and the Sensor-1 residue T/N and the basic arginine residues from the Sensor 2 helix also H-bond with the phosphate group of ATPγS (**Fig. 3a-c**). The adjacent subunit also contributes to nucleotide binding. Unexpectedly, the two arginine residues from the central α5 helix and the SRC-containing α6 helix each form H-bonds with the γ-phosphate in the Rad24-RFC–9-1-1–DNA structures, indicating that the first four nucleotide binding sites are poised for ATP hydrolysis, even though the Rad24-RFC is nearly planar (**Fig. 3b**). This is different from the spiral RFC–PCNA structure in which only the ATP analogue in the first interface between RFC1 and RFC2 is tightly bound. In the following three ATP sites, the arginine finger residues of the SRC motifs are 8-17 Å away from the γ-phosphate, indicating an inactive ATPase state (Gaubitz et al., 2020). This difference between the nearly planar Rad24-RFC and spiral RFC structures is likely due to the presence of DNA in the Rad24-RFC structure, as DNA binding and the subsequent clamp loading likely induce conformation changes that could switch the loader from the inactive to active state, eventually triggering ATP hydrolysis and the release of the loaded DNA clamp from the clamp loader (Bowman et al., 2004; Gaubitz et al., 2020; Kelch et al., 2011)

**Fig 3.**
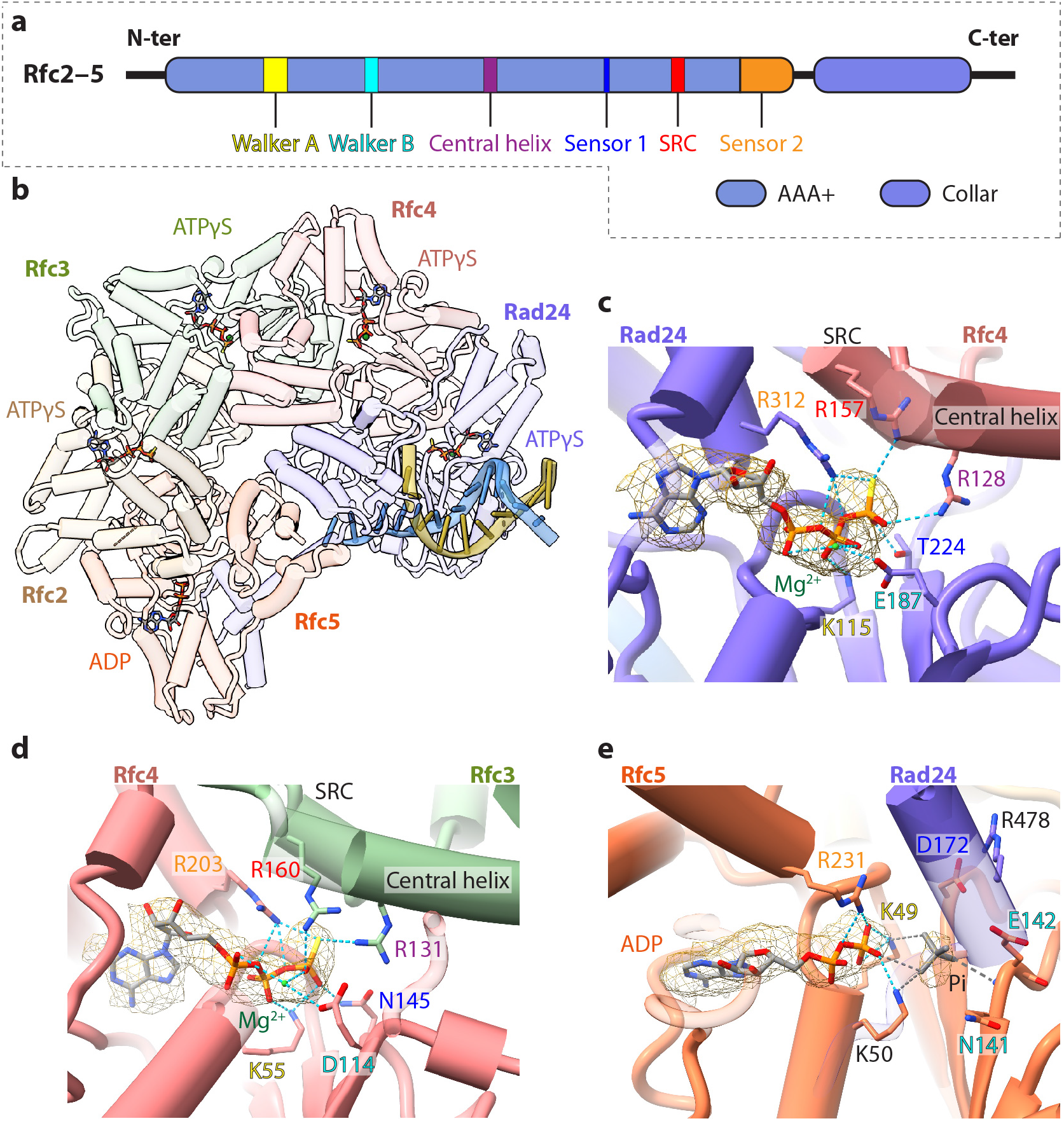
Four ATPγS and one ADP are bound to Rad24-RFC. **a**) Key ATPase motifs in Rfc2-5. Rad24 lacks the SRC and central helix. **b**) Top view of Rad24-RFC–DNA in cartoon, omitting the 9-1-1 clamp for clarity. The bound four ATPγS and one ADP are shown in sticks. **c-e**) Enlarged views of the nucleotide binding pockets in: c) Rad24, d) Rfc4, and e) Rfc5. ATPγS binding in Rfc2 and Rfc3 are similar to that in Rfc4 and, therefore are not shown. The residues involved in nucleotide binding in the conserved SRC (serine-arginine-cysteine motif), sensor-1, Walker A (P-loop), Walker B (DExx box), Sensor-2, and the central helix are labeled and colored red, orange, yellow, cyan, blue, and purple, respectively. Mg^2+^ ion is green. The Rfc5 nucleotide pocket is occupied by ADP. Because Rad24 lacks the SRC motif, there is only one arginine (Arg-478) at the Rfc5:Rad24 interface, which points away from the nucleotide due to the absence of the γ-phosphate. The isolated strong density (displayed at 6σ) next to ADP is modeled as a phosphate (P_i_) and shown in grey sticks. The phosphate is coordinated by both Lys-49 and Lys-50 of the Walker A motif as well as the backbone nitrogen of Glu-142 in the Walker B motif. For clarity, only the hydrogen bonds with the nucleotide phosphates are shown.

The presence of ADP at the interface of Rfc5:Rad24 is remarkable, as this resembles those observed at the equivalent site in all three structurally determined clamp–clamp loader systems, despite the significant differences between Rad24 and Rfc1 (Bowman et al., 2004; Gaubitz et al., 2020; Kelch et al., 2011) (**Fig. 3d**). Rfc5 harbors all essential ATPase motifs. However, at the opposing side of the interface, the Rad24 C-terminal A’ subdomain has only one helix (α-18) that harbors an arginine residue (Arg-478), as compared to two helices – the SRC-containing helix and the central helix at all other ATP binding interfaces. The Walker A motif of Rfc5 – ^43^<u>G</u>PN<u>G</u>T<u>GK</u>*K*T^51^ – has an extra lysine (Lys-50) (Cai et al., 1998; Gaubitz et al., 2020; Venclovas et al., 2002) (**Extended Data Fig. 7**). A previous study found that mutation of the first lysine (Lys-49) alters neither the ATPase activity nor DNA replication activity of the RFC complex, leading to the suggestion that this nucleotide binding site is inactive in ATP hydrolysis, and that the observed ADP may be copurified with the loader or as a contaminant that is typically found in ATPγS preparations (Cai et al., 1998; Gaubitz et al., 2020). Interestingly, the Rfc5 Lys-50 is involved in ADP binding in our structure, potentially explaining the mutation tolerance of Lys-49, because of the compensating effect of Lys-50. More importantly, there is a bulky density near the ADP β-phosphate in our EM map when displayed at a high threshold of 6σ (**Fig. 3d**). This density is adequate to accommodate a phosphate and can be properly coordinated by surrounding residues. Therefore, the observed ADP and phosphate in our Rad24-RFC structure are likely the hydrolytic products of ATPγS. Based on our structure, we suggest that the extra lysine (Lys-50) in the Walker A motif of Rfc5 may have enhanced the ATPase activity at this nucleotide binding interface, such that the ATPγS bound there has been hydrolyzed to ADP and γS-phosphate. In this scenario, the released phosphate may have diffused into solvent in the previous clamp loader structures (Bowman et al., 2004; Gaubitz et al., 2020; Kelch et al., 2011).

### 4. Interactions between Rad24-RFC and the 9-1-1 clamp

Rad24-RFC features three insertion loops that are important to its binding to the DNA substrate as well as the 9-1-1-clamp (**Fig. 4a**). The first loop is a long insertion in the AAA+ domain of Rad24, termed the upper loop. The second loop is an insertion in Rfc5, which we call the Rfc5 plug, that is much longer than those of Rfc2-4 (**Extended Data Fig. 7**). These two loops project into the interior chamber of the Rad24-RFC and constrain the DNA path to 12 Å, which is narrower than the 20-Å dsDNA width. Thus, these two loops may prevent dsDNA binding to the inner chamber of Rad24-RFC and may facilitate the inverted DNA binding mode in Rad24-RFC, as compared to the RFC in which the interior chamber is wide enough to accommodate the dsDNA. The third loop is hook-like, inserted in the bottom surface of the Rad24 AAA+ domain and interacts with the Rad17 subunit of the yeast 9-1-1 clamp in the closed state. This loop is ordered in the clamp closed state but becomes disordered in the clamp open state (**Fig. 4a-b**). This clamp binding feature is found only in the Rad24 family of proteins and is absent in the RFC complexes (**Extended Data Figs. 5, 7**).

**Fig. 4.**
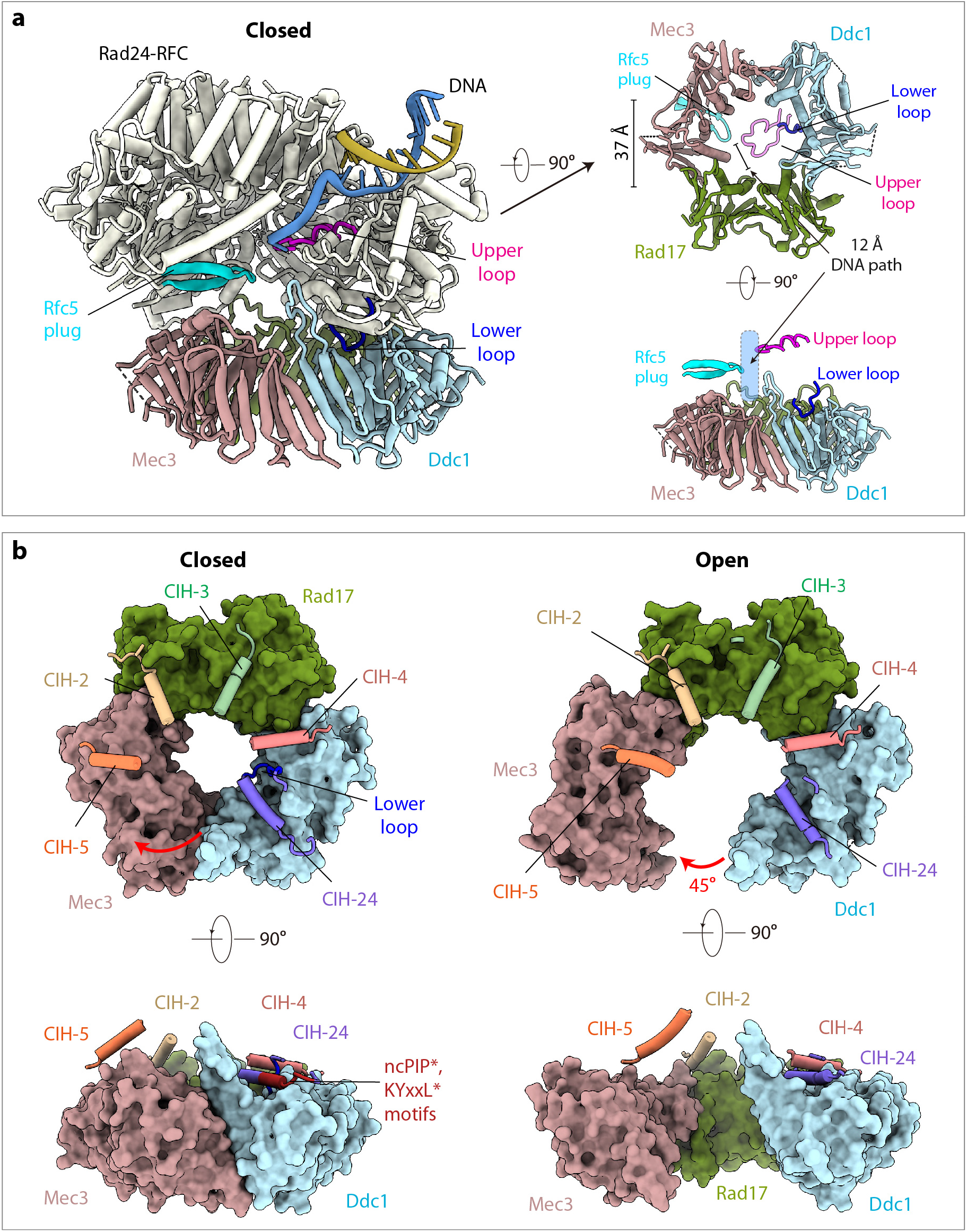
Interaction between Rad24-RFC and the 9-1-1 clamp in the closed and open state. **a**) Tilted, bottom, and side views of the closed 9-1-1 clamp, highlighting the positions of three important loops. The upper insertion loop of Rad24 and the Rfc5 plug narrow the interior chamber of Rad24-RFC to 12 Å, suitable for ssDNA but excluding dsDNA binding. The Rad24 lower insertion loop binds ddc1 of the 9-1-1 clamp to control the ring opening/closure. The inner diameter of 9-1-1 is 37 Å. **b**) Comparison of the closed and open 9-1-1 clamp in top and side surface views. Superimposed on the clamps (in surface view) are the clamp-interacting helix (CIH) and connecting loop of each loader subunit shown in cartoon view. The noncanonical PIP (ncPIP) motif and the KYxxL-like motif at the end of the Rad24 CIH are labeled and highlighted in dark red. The Rad24 lower loop bound to Ddc1 is ordered in the closed state but becomes disordered and loses contact with Ddc1 in the open state, leading to a 45° in-plane rotation of Mec3 to form the DNA entry gate.

Interactions between Rad24-RFC and 9-1-1 are primarily mediated by the clamp interacting α4-helices (CIHs) and the loops that follow them, and these interactions are similar to those between RFC and PCNA (Bowman et al., 2004; Gaubitz et al., 2020) (**Fig. 4a-b, Extended Data Figs. 5, 7**). However, only 3 RFC subunits were bound to the PCNA ring in the human RFC–PCNA structure, because the PCNA ring and the RFC spiral are at an angle. In contrast, all five subunits of Rad24-RFC had some contacts with the 9-1-1 clamp in both closed and open states, due to the planar structure of both complexes and thus the close approach of the 9-1-1 ring to the loader such that the 9-1-1 ring and the Rad24-RFC ring are nearly parallel to each other. In the closed conformation, Rad24 sits above Ddc1, having the largest interface of 830 Å^2^. But this interface is weakened and reduced by one third in the open state. The interface between Rfc2 and Ddc1 is 400 Å^2^ in the closed state and is reduced slightly to 350 Å^2^ in the open state. The interfaces between Rad17 and Rfc3 and between Rad17 and Rfc4 are stable, with a total area of 940 Å^2^ in both closed and open states. Compared to the buried surface of 2,000 Å^2^, 700 Å^2^ and 1,200 Å^2^ between PCNA and RFC1, RFC2 and RFC3, respectively (Gaubitz et al., 2020), the binding between Rad24-RFC and 9-1-1 is substantially weaker. However, it should be noted that the binary RFC–PCNA complex was assembled in the absence of DNA, and the complex was stabilized by chemical crosslinking (Gaubitz et al., 2020). Thus, it is unclear if a native (uncross-linked) RFC–PCNA complex maintains such an extensive interface.

Overall, Rad24 contributes the most binding affinity for the 9-1-1 clamp. Rad24 lacks the canonical PCNA-interacting peptide (PIP) motif of <u>Q</u>xx<u>Ψ</u>xx<u>θθ</u>, where Ψ is hydrophobic, θ is aromatic, and x is any residue (Prestel et al., 2019). However, based on sequence alignment, we identified a non-canonical PIP motif ^169^<u>F</u>LK<u>G</u>AR<u>YL</u>^176^ in the AAA+ domain, and this motif interacts with the 9-1-1 clamp (**Fig. 4b, Extended Data Fig. 5**). Interestingly, the last three residues of the non-canonical Rad24 PIP motif plus the following two residues (^174^<u>RY</u>LV<u>M</u>^178^) are equivalent to the <u>KY</u>xx<u>L</u> motif in the human Rad17 that was previously shown to be essential for 9-1-1 binding (Fukumoto et al., 2016); indeed, this 5-residue motif also contributes to the yeast 9-1-1-clamp binding in our structure.

### 4. Both the open and closed 9-1-1 clamps are planar

The Rad24-RFC loader is configured into a two-tiered spiral, with the upper tier formed by the helical “collar” domain and the lower tier by the AAA+ ATPase domains that engages the clamp ring (**Figs. 1d, 4a**). The overall size of the Rad24-RFC–9-1-1–DNA complex is similar in both states, with the closed state (132 Å × 121 Å × 116 Å) being more compact than the open state (134 Å × 118 Å × 114 Å) (**Extended Data Fig. 3a-b**). The structures of the two conformations are also similar with an RMSD of 1.0 Å when the two gap-lining 9-1-1 subunits Ddc1 and Rad17 are excluded from comparison (**Fig. 4b**). We found that Mec3 of the closed form undergoes a 45° in-plane rotation away from Ddc1 to generate the 27-Å gap in the open form of the 9-1-1 clamp (**Supplementary Video 1**). This is the largest gap observed so far in any DNA clamp-loader complex structure; the previously observed open clamp rings only has a gap of 10 Å or less that is too narrow for dsDNA to pass through (Bowman et al., 2004; Gaubitz et al., 2020; Kelch et al., 2011; Liu, 2019; Miyata et al., 2005). Importantly, the gate opening coincides with the above-mentioned disordering and perhaps unbinding of the Rad24 lower insertion loop (**Fig. 4b**). Therefore, we suggest that this Rad24 loop controls the 9-1-1 ring opening.

The pre-existing RFC–PCNA structures were determined in the absence of a DNA substrate in which the PCNA is a closed ring and the RFC loader is spiral (Bowman et al., 2004; Gaubitz et al., 2020). Compared with the RFC-PCNA structure, the 9-1-1 clamp tilts up by 25° to engage the loader in our DNA-bound Rad24-RFC–9-1-1 structure (**Extended Data Fig. 8, Supplemental Video 1**). Furthermore, the Rad24-RFC loader is wider than the RFC loader, due to a large expansion of the Rad24 structure to accommodate the DNA. However, the close approach of the 9-1-1 ring towards Rad24-RFC resembles the DNA-bound T4 phage clamp–clamp loader structure (Kelch et al., 2011). Strikingly, the 9-1-1 ring is planar in both closed and open states of the Rad24-RFC–9-1-1 clamp–DNA complex. This contrasts with the spiral T4 clamp loader-clamp with DNA (Kelch et al., 2011). While the physiological role of maintaining a planar 9-1-1 ring during the DNA loading process is currently unclear, we speculate this unique feature may be related to the fact that 9-1-1 is loaded onto ssDNA, not dsDNA, and then slides to the 5’ dsDNA/ssDNA junction after it is loaded and separates from the Rad24-RFC complex.

### 6. Proposed 9-1-1 clamp loading mechanism by Rad24-RFC

The ssDNA binding protein RPA is known to be involved in Rad24-RFC function and its specificity for the 5’-junction (Majka et al., 2006). This function is mediated by a protein-protein interaction between the 180-residue N-terminal domain of Rpa1 and the C-terminal coiled coil domain (aa 473 – 652) of Rad24 (Majka et al., 2006; Piya et al., 2015). The Rad24 C-terminal coiled coil is disordered in our structure determined in the absence of RPA. But its approximate location can be predicted following the visualized A’ subdomain to which the coiled coil is appended. Based on our structure, we suggest that RPA–Rad24-RFC together provide a bipartite damaged DNA binding platform, with Rad24-RFC binding at the 5’-junction and the RPA binding to the ssDNA before the junction, which is a 3’ overhang of ssDNA prior to the 5’ ss/ds DNA junction (**Fig. 5a**, left panel). Such bipartite binding likely holds the 3’-overhang ssDNA away from the space below the AAA+ ring of Rad24-RFC, thereby clearing the way for 9-1-1 to approach and engage the Rad24-RFC on the 5’-DNA junction. We speculate that RPA may further align the 3’-overhang ssDNA from the 5’-DNA junction with the open gate of the 9-1-1 clamp at the subsequent 9-1-1 clamp loading stage, in a manner akin to Orc5-6 of the ORC complex holding one end of the replication origin DNA away from the Mcm2-7 binding space and aligning the DNA with the Mcm2-Mcm5 DNA gate of the Cdt1-bound Mcm2-7 (Yuan and Li, 2020; Yuan et al., 2017; Yuan et al., 2020).

**Figure 5.**
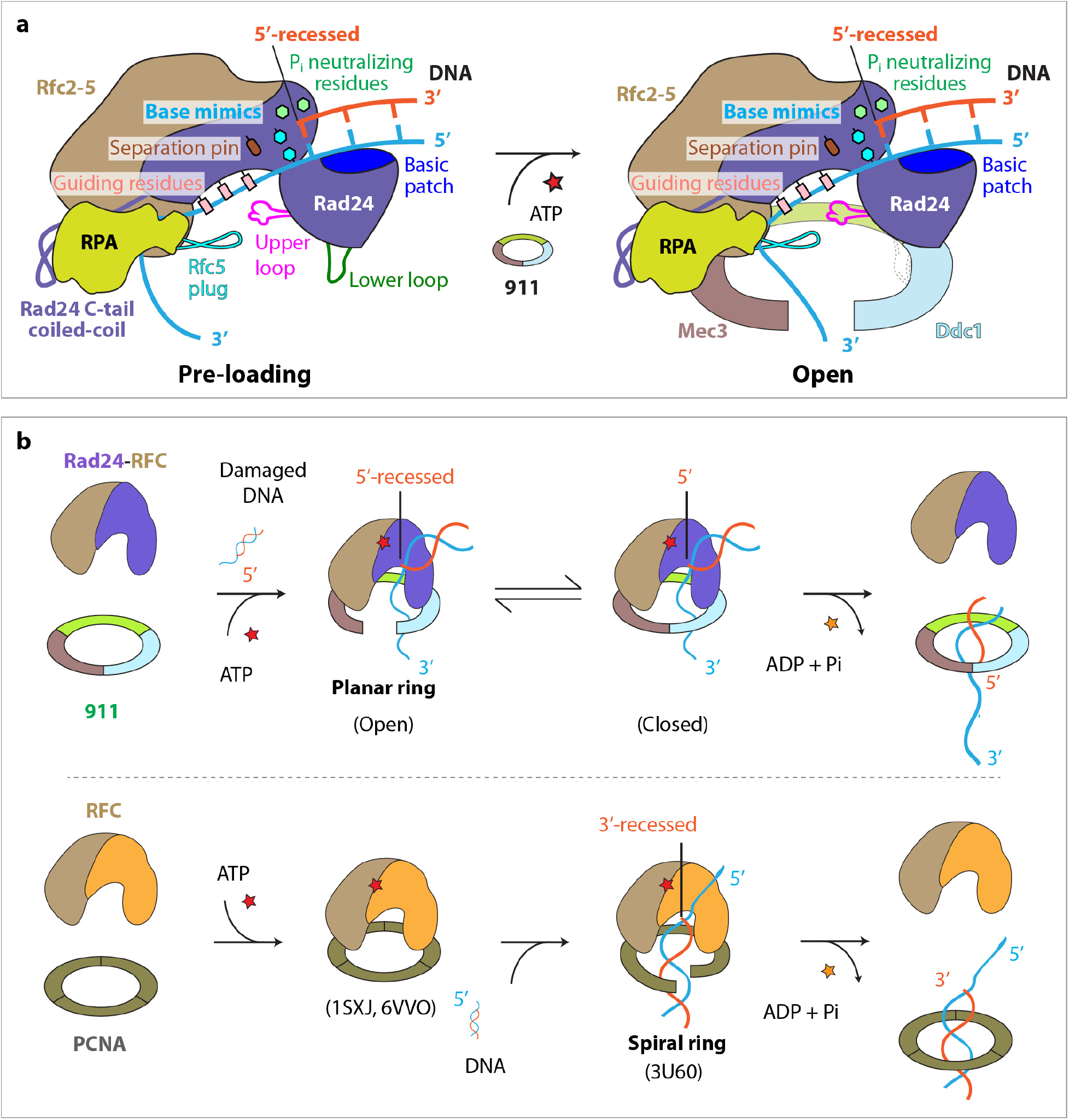
The mechanism of 9-1-1 loading onto the 5’-DNA junction by Rad24-RFC. **a**) Rad24 confers the 5’-DNA junction specificity via several unique structural features: the long linker between the collar and AAA+ domains enabling their separation and the formation of the dsDNA binding groove; the acidic patch on top of the AAA domain that binds dsDNA; the 5’-phosphate neutralizing residues, the base mimics, and the separation pin that facilitate binding to the 5’-junction; and the Rad24 upper loop and the Rfc5 plug that limit the chamber size and prevent dsDNA from entering the loader chamber. RPA binds to the C-terminal coiled coil of Rad24, thereby holding the ssDNA of the 5’-DNA junction away from the space below the Rad24-RFC where the 9-1-1 clamp will bind. This is a bipartite DNA binding mechanism that facilitate 9-1-1 clamp recruitment and loading onto ssDNA at a 5’ DNA junction. The 911 clamp could then diffuse onto the dsDNA **b**) Upper panel: The loading steps of the 9-1-1 clamp by Rad24-RFC. Both open and closed states are observed in this study. Lower panel: the loading steps of PCNA by RFC based on previous structural studies. In addition to the opposite DNA binding mode, the inner chamber of RFC is a spiral that is molded by the binding of DNA, whereas Rad24-RFC does not adopt a spiral structure as the dsDNA region is solely bound by the Rad24 subunit and does not enter the inner chamber of Rad24-RFC. See text for details.

While RPA appears to be the first responder to DNA damage, it acts together with Rad24-RFC to mount an effective response. We suggest that RPA holds the 3’-overhang ssDNA at a 5’-DNA junction, Rad24-RFC specifically recognizes the 5’-junction and binds the dsDNA region between the collar domain and AAA+ domain of Rad24. The 5’-junction recognition is likely mediated by a number of unique Rad24 residues, including the three residues that neutralize the 5’-phosphates (e.g. His-341-, Lys-345, and His-351), the DNA separation pin at the junction (Tyr-442), and the two base-mimicking residues (e.g, Phe-340 and His-341) that may stabilize the last base pair at the junction (**Fig. 5a**, left panel). Next, Rad24-RFC recruits the 9-1-1 clamp to the DNA damage site. The DNA-bound Rad24-RFC first binds the 9-1-1 via the five clamp-interacting helices (CIH), one from each loader subunit, and then inserts a hook-like loop at the bottom surface of the Rad24 AAA+ domain into the 9-1-1 clamp and binds exclusively to the ddc1 clamp subunit (**Fig. 4b**, left panel). This binding event on Ddc1 likely triggers the neighboring Mec3 clamp subunit to undergo an in-plane rotation away from Ddc1, generating a gap wide enough to admit either a ssDNA or a dsDNA. Loading of the 9-1-1 on the DNA will presumably stimulate ATP hydrolysis by Rad24-RFC, leading to ring closure of the 9-1-1 clamp and its separation from the Rad24-RFC. The ATP hydrolysis coupled separation mechanism is supported by the structural analysis of the T4 clamp loader bound to a DNA and a closed clamp ring, which revealed that ATP hydrolysis at the subunit B:C interface results in subunit B to swing away from subunit C, releasing subunit B from the clamp (**Extended Data Fig. 8**) (Kelch et al., 2011). Departure of the clamp loader leads to the exposure of the 9-1-1 ring for recruitment of downstream checkpoint proteins to the damage site for DNA repair, such as the ATR kinase (yeast Mec1-Ddc2). This contrasts with the RFC-PCNA system in which departure of the RFC loader facilitates the recruitment of enzymes such as the replicative polymerases δ and ε (Kelch et al., 2012).

The mechanism of PCNA loading on DNA by the eukaryotic RFC has been studied extensively (Kelch et al., 2012; O’Donnell and Kuriyan, 2006). Structural determination of the RFC-PCNA in the absence of a DNA substrate has led to the proposal that RFC first engages PCNA, and then the RFC-PCNA binary complex engages the 3’-DNA junction, leading to PCNA loading and encircling the dsDNA (Bowman et al., 2004; Gaubitz et al., 2020) (**Fig. 5b**, lower panel). There is one reported structure of the clamp–clamp loader complex bound to a DNA substrate – that of the T4 phage system (Kelch et al., 2011). In that structure, the 3’-DNA junction has entered the inner collar region, but the 5’-overhang is extruded from the chamber (**Extended Data Fig. 8a** and **c**). That structure may represent the post-loading state. A larger gate opening may have been available to admit the dsDNA but the gate has been partially closed once the DNA has entered. Another possibility is that clamp may initially be loaded onto ssDNA and then the 3’ ss/ds junction may slide into the clamp, which would not require a ring opening larger than that needed to accommodate ssDNA (Bowman et al., 2004; Kelch et al., 2011).

In sum, the work described here provides the first structural support for a highly unusual and unexpected mechanism for the Rad24-RFC specificity for a 5’-DNA junction. Our structures reveal that Rad24-RFC binds the ds DNA of a 5’-DNA junction above the plane of an open ring, in which both the ring and the clamp loader are essentially in parallel to each other. Furthermore, the Rad24-RFC directs ssDNA into the 9-1-1 clamp, rather than dsDNA. To accommodate this drastic difference between RFC and Rad24-RFC, there are several Rad24-specific features to load the 9-1-1 clamp in an orientation that is opposite of the RFC loading of the PCNA clamp.

## Acknowledgements

Cryo-EM micrographs were collected at the David Van Andel Advanced Cryo-Electron Microscopy Suite in Van Andel Research Institute, Grand Rapids, MI. We thank G. Zhao and X. Meng for facilitating data collection. This work was supported by the US National Institutes of Health grants GM115809 (to M.E.O.) and GM131754 (to H.L.), A BCRF 20-068 grant (o M.E.O,), Howard Hughes Medical Institute (to M.E.O), and the Van Andel Institute (to H.L.).

## Author contributions

H.L. and M.E.O. designed research; F.Z., N.Y. and R.G. performed research; F.Z., N.Y., R.G., M.E.O. and H.L. analyzed the data and wrote the manuscript with input from all authors.

## Competing interests

The authors declare no competing interests.

## Data Availability

The 3D cryo-EM maps of S.c. Rad24-RFC–9-1-1 clamp-DNA complexes in the closed and open conformation at 3.2-Å resolution have been deposited in the Electron Microscopy Data Bank with accession codes EMD-xxxx1 and EMD-xxxx2. The corresponding atomic models have been deposited in the Protein Data Bank with accession codes xxx1 and xxx2.

## MATERIALS AND METHODS

### Cloning and expression of Rad24-RFC

The alternative clamp loader Rad24-RFC was expressed in *E. coli* by cloning the five genes into two compatible expression vectors that have different bacterial plasmid origins. The Rfc2, 3 and 4 genes were cloned into pET3c (ColE1 origin) under control of the T7 promotor (Amp selection). The Rad24 and Rfc5 genes were cloned into the pLANT vector (p15a origin), under control of the T7 vector (Kan selection). We have previously described the pLANT expression vector (Finkelstein et al., 2003). To express the Rad24-RFC, the pET-RFC24/Rfc5 and pLANT Rfc2, Rfc3, Rfc4 plasmids were cotransformed into competent *E. coli* BL21(DE3) codon plus cells which require Cam to maintain the codon plus plasmid that carries tRNAs common to eukaryotes but uncommon to E. coli (Invitrogen). The transformed cells were plated on LB containing 100 μg/ml Amp, 50 μg/ml Kan and 25 μg/ml Cam. A single colony was used to inoculate a 125 ml culture in LB containing these same antibiotics. After shaking at 37°C for 3 h, 10 ml of cell culture was added to each of 12 2-L fluted flasks that each contained 1L of LB containing the same antibiotics. The flasks were incubated in a shaker floor incubator for 19 h at 30°C at which time cultures reached an OD_600_ of 0.6. Cell cultures were then brought to 15°C by swirling each flask in ice water using a thermometer to gauge the temperature. Flasks were then placed in a different floor shaker that had been pre-equilibrated at 15°C. Expression was then induced upon adding 1 mM IPTG. Cells were allowed to express for 12h, then were harvested by centrifugation and resuspended in 180 ml Hepes pH 7.5, 1 mM DTT, 0.5 mM EDTA, 20% glycerol, 400 mM NaCl.

### Purification of Rad24-RFC

Cells were lysed by French Press using 3 passes at 22,000 psi each pass. The lysate was clarified by centrifugation at 12,000 rpm in a Sorvall SLA 1500 rotor for 1 h at 4°C. Clarified lysate was diluted with buffer A (Hepes pH 7.5, 1 mM DTT, 0.5 mM EDTA, 20% glycerol) to a conductivity equal to 150 mM NaCl, and then loaded onto a 125 ml SP Sepharose column equilibrated in buffer A containing 150 mM NaCl. The column was washed with buffer A + 150 mM NaCl until the UV_280_ reached a baseline level, and then elution was performed with a 1-liter linear gradient from 150 mM NaCl to 600 mM NaCl in buffer A. Fractions of 15 ml were collected and analyzed for Rad24-RFC by 10% SDS PAGE. Fractions 16-22 (140 ml) were pooled. The pool was diluted 1.5-fold using buffer A to a conductivity of 200 mM NaCl, and then loaded onto a 50 ml Fast Flow Q column pre-equilibrated in buffer A + 150 mM NaCl. The column was washed with buffer A + 150 mM NaCl until the UV_280_ approached background, and then was eluted with a 500 ml linear gradient of buffer A from 150 mM NaCl to 600 mM NaCl, collecting 7.5 ml fractions. Fractions 12-18 (120 mg) were pooled and aliquoted into 1 ml amounts and stored at −80°C.

### Cloning and expression of the 9-1-1 complex

Rad17, Ddc1 and His-PK-Mec3 (Mec3 containing a N-terminal hexa-histidine tag, plus a 7-residue protein kinase A recognition site (LRRASLG) (Kelman et al., 1995), were cloned onto pET21a for expression by T7 RNA polymerase. The pET-911 plasmid was transformed into BL21(DE3) codon plus cells (Invitrogen) and plated on LB plus 100 μg Amp, 25 μg Can. A single colony was used to inoculate 125 ml LB containing 100 μg Amp, 25 μg at 37°C with shaking for 2 h. Then 10 ml of the cell suspension was added to each of 12 2 L fluted flasks that each contained 1 L LB containing 100 μg Amp, 25 μg Can. The cell cultures were grown to OD_600_ and then each flask was swirled in ice water to bring cultures to 15°C. The flasks were then transferred to a precooled floor shaker incubator at 15°C and IPTG was added to 1mM. Cells were induced for 10 h at 15°C prior to harvesting by centrifugation. Cells were resuspended in an equal volume (to the weight of the cell pellet – via weighing centrifuge bottles before and after cell collection) of 20 mM Tris-Cl pH 7.5, 10% w/v sucrose and frozen at −80°C.

### Purification of the 9-1-1 complex

Cell pellets equal to 6 liters of cell expression culture, were thawed and brought to a volume of 75 ml using buffer B (25 mM Tris-Cl pH 7.5, 1 mM DTT, 0.5 mM EDTA, 10% glycerol). Cells were then lysed by three passages through a French Press operating at 22,000 psi. The lysate was clarified by centrifugation at 12,000 rpm in a Sorvall SLA 1500 rotor for 1 h at 4°C. Clarified cell lysate (240 ml) was treated with 96 g (NH4)2SO4 with slow stirring for 30 min, then the protein precipitate was collected by centrifugation in an SLA rotor at 12,500 rpm for 30 min at 4°C. The supernatant was discarded, and the pellet was resuspended in 24 ml of Ni column binding buffer (20 mM Tris-Cl pH 7.9, 5 mM Imidazole, 0.5 M NaCl, 10% glycerol) and dialyzed against 1L Ni column binding buffer overnight in the cold room. Protein was then loaded onto a 20 ml chelating Nickel HiTrap column (Sigma) that had been equilibrated in Ni column binding buffer. The HiTrap column was washed with 40 ml Ni column binding buffer + 60 mM imidazole, and then eluted with a 200 ml linear gradient of 60 mM – 1M imidazole in Ni column binding buffer. Fractions of 2 ml were collected and then analyzed by SDS PAGE. Fractions containing 9-1-1 were pooled, dialyzed against buffer B containing 150 mM NaCl overnight, and then loaded onto a 40 ml SP Sepharose column equilibrated in Buffer B + 150 mM NaCl. The column was washed with 100 ml Buffer B + 150 mM NaCl and then eluted with a 400 ml linear 150 mM – 600 mM NaCl in Buffer B. Fractions of 5 ml were collected and analyzed by SDS-PAGE. The 9-1-1 flows through the column at this ionic strength, but many contaminants stuck to the column and were thus removed. The flow through, containing 9-1-1, was then dialyzed against Ni column loading buffer and loaded onto a second Ni column of 5-ml bed volume. The loaded column was washed with 10 ml Ni column binding buffer + 60 mM imidazole, and then eluted with a 50 ml gradient of 60 mM – 1M imidazole in Ni column binding buffer. Fractions of 1 ml were collected and analyzed by SDS PAGE. Fractions containing 9-1-1 were pooled, and the nearly pure 911 was dialyzed against Buffer B containing 50 mM NaCl. The protein was then loaded onto a 1 ml MonoQ column and eluted with a 10 ml linear gradient from 50 – 500 mM NaCl in Buffer B. Fractions of 0.25 ml were collected and analyzed by SDS PAGE. Fractions 29-33 contained pure 9-1-1 (approximately 2 mg) and were aliquoted and stored at −80°C.

### Cryo-EM grids preparation and data collection

The 3’-junction DNA substrate was composed of the template strand (5’-CTG CAC GAA TTA AGC AAT TCG TAA TCA TGG TCA TAG CT-3’, 38 nt) and the primer strand (5’-AGC TAT GAC CAT GAT TAC GAA TTG-ddC-3’, 25 nt). The DNA with both 3’- and 5’-junctions (double tailed DNA) was composed of Watson strand (5’- GCA GAC ACT ACG AGT ACA TA-3’, 20 nt) and Crick strand (5’- TTT TTT TTT TTA TGT ACT CGT AGT GTC TGC TTT TTT TTT T-3’, 40 nt). These DNAs were synthesized annealed, and then HPLC purified by Integrated DNA Technologies, Inc. The assembly of yeast Rad24-RFC–9-1-1 clamp–DNA complex followed our previous procedure for assembling the S.c. Pol δ–PCNA-DNA complex in vitro (Zheng et al., 2020). Briefly, a droplet of 10 μl of purified 9-1-1 clamp at 3.3 μM and 0.4 μl of 3’-junction DNA or the double tailed DNA at 100 μM were mixed and incubated at 30°C for 10 min, then the mixture and 0.75 μl 10 mM ATPγS were added into 3.2 μl of purified Rad24-RFC protein at 8.6 μM concentration. The final concentration of the reaction mixture is 1.8 μM Rad24-RFC, 2.2 μM 9-1-1 clamp, 2.7 μM DNA (either the 3’-junction DNA or the double tailed DNA), 0.5 mM ATPγS, and 5 mM MgAcetate, with a total reaction volume of 15-μl. The mixture was then incubated in an ice-water bath for additional 3 hr (3’-junction DNA) or 30 min (double tailed DNA). The final molar ratio of Rad24-RFC: 9-1-1 clamp: DNA was 1.0: 1.2: 1.5 with both DNA substrates. The Quantifoil Au R2/1 300 mesh grids were glow-discharged for 1 min in a Gatan Solarus, then 3 μl of the mixture was applied onto the EM grids. Sample vitrification was carried out in a Thermo Fisher Vitrobot Mark IV with the following settings: blot time 2 s, blot force 4, wait time 1 s, inner chamber temperature 6°C, and a 95% relative humidity. The EM grids were flash-frozen in liquid ethane cooled by liquid nitrogen. Cryo-EM data were automatically collected on a 300 kV Titian Krios electron microscope controlled by SerialEM in a multi-hole mode. The micrographs were captured at a scope magnification of 105,000×, with the objective lens under-focus values ranging from 1.1 to 1.9 μm, by a K3 direct electron detector (Gatan) operated in the super-resolution video mode. During a 1.5 s exposure time, a total of 75 frames were recorded with a total dose of 65 e^-^/Å^2^. The calibrated physical pixel size was 0.828 Å for all digital micrographs.

### Image processing and 3D reconstruction

For the double-tailed DNA bound ternary complex, the data collection and image quality were monitored by the cryoSPARC Live v3.2 installed in a local workstation (Punjani et al., 2017). The image preprocessing including patch motioncorrection, CTF (contrast transfer function) estimation and correction, blob particle picking (70-150 Å diameter) and extraction with a binning factor of 2 were also carried out at the same time, and a total number of 14,521 raw micrographs were recorded during a three-day real-time data collecting session. About 8.5 million particles were extracted and subjected to two rounds of 2-dimentional (2D) image classification, which resulted in ~471,000 “good” particle images. These particles were used to calculate four starting 3D models. At this stage, we observed two good 3D classes representing the Rad24-RFC–9-1-1 clamp–DNA complexes in clamp-closed and clamp-open conformations. Next, we used Topaz (including training and picking) to pick more particles or particles in more views (Bepler et al., 2019). To avoid missing those less frequently occurring particle views, we used the recently reported “Build and Retrieve” method to further clean up the particle images extracted by Topaz (Su et al., 2021). Finally, the cleaned-up particles were combined, and duplicate particles with 40% or larger particle diameter overlapping (56 Å) were removed. After homogenous and non-uniform 3D refinement, we performed 3D viability analysis (3DVA) on 3 3D EM modes in 30 iterations. This classified the particles into five 3D classes. A gate-closed conformation with approximately 147,000 particles and an open-gate conformation with approximately 300,000 particles were chosen for further non-uniform, per-particle CTF, and local refinement, yielding the two final 3D maps at an average resolution of 3.17 Å (closed) and 3.23 Å (open), respectively. The 9-1-1 clamp regions in the two final 3D maps were more flexible and had weaker densities than the upper Rad24-RFC loader region. We performed 3D focused refinement and obtained the clamp densities to 3.8 Å resolution in the gate-closed state and to 4.3 Å resolution in the gate-open state, respectively. The improvement of the focused refinement was marginal. Thus, we used the two final full maps (without focused refinement) for model building.

For the sample mixed with the 3’-junction DNA, we recorded 12,000 raw micrographs under similar data collection procedures. After 2D image classification, about 230,000 particles were retained. Based on 2D and 3D classifications, we found that neither a binary complex of Rad24-RFC–9-1-1 clamp nor a ternary complex of Rad24-RFC–9-1-1 clamp–DNA formed in the presence of the 3’-junction DNA.

### Model building, refinement, and validation

For the atomic model building of the *S. cerevisiae* Rad24-RFC–9-1-1 clamp–DNA complex 3D maps in the closed and open states, we used the two homologous crystal structures of the human 9-1-1 clamp structure (PDB code 3G65) and the yeast RFC–PCNA structure (PDB code 1SXJ) as initial models for the yeast 9-1-1 and Rfc2-5 respectively. Because no Rad24 or its human homologue Rad17 structures were available, a *de novo* cryo-EM map modelling server DeepTracer (https://deeptracer.uw.edu) was used to produce the starting model (Pfab et al., 2021). The DNA density in our 3D EM map is sufficient to distinguish a purine from a pyrimidine. We used the *de novo* modeling program Map-to-Model wrapped in PHENIX (Adams et al., 2010) to obtain the initial DNA model.

Although there are several human 9-1-1 clamp structures available (Dore et al., 2009; Sohn and Cho, 2009; Xu et al., 2009). Model building for the yeast 9-1-1 ring was more challenging, because the map had lower resolution (3.8 Å) in this region (**Extended Data Fig. 3**), and the three subunits (Ddc1, 612 aa; Mec3, 474 aa; Rad17, 401 aa) are divergent from their respective human analogues (Rad9, 391 aa; Hus1, 280 aa; Rad1, 282 aa). There are several long insertions in each yeast subunit compared to the structurally conserved PCNA-like core (Venclovas and Thelen, 2000). Thus, we adopted a homologue modelling approach using the human 9-1-1 clamp structure as a template. We first used the Phyre2 server (http://www.sbg.bio.ic.ac.uk/phyre2) (Kelley et al., 2015) to predict the structures of the three subunits. We next used I-TASSER (Yang et al., 2015) and Robetta (Song et al., 2013) to predict these subunits independently. The models predicted by these three programs were very similar, and we chose the predictions from Phyre2 as the starting models. We only kept the PCNA-like core region and removed the peripheral insertion loops, because these loops are flexible and invisible in our EM maps. We then performed flexible fitting into the EM maps using the Flex-EM program wrapped in the CCP-EM package (Burnley et al., 2017; Topf et al., 2008).

Finally, the flexibly fitted 9-1-1 ring, the *de novo* built Rad24 and DNA models, and the yeast RFC-PCNA structure were fitted together into the EM map in the closed conformation. Then, the PCNA and Rfc1 were removed, and the remaining models were merged into a single file serving as the starting model of the complex in the UCSF Chimera (Pettersen et al., 2004). This starting model was refined iteratively between the real space computational refinement in PHENIX and manual adjustment in COOT (Emsley et al., 2010). The final model of the closed conformation was refined to 3.2 Å and went through a comprehensive validation by the MolProbity program embedded in PHENIX (Chen et al., 2010). The closed conformation model was used as the initial model for the open state EM map. The model was iteratively refined and manually adjusted as described above, except for the 9-1-1 subunit Ddc1 and Mec3, which had much weaker EM densities; these two subunits were modelled by rigid body docking into the EM densities. Structure figures were prepared using ChimeraX (Pettersen et al., 2021) and organized in Adobe Illustrator (Adobe Inc).

## SUPPLEMENTARY MATERIALS

**Supplementary Table 1.** Cryo-EM data collection, refinement, and atomic model validation. The first 7 parameters apply to both open and closed states, as depicted in the workflow of Extended Data Fig. 2.

| **Rad24-RFC-9–1-1 clamp–DNA ternary complex** | **Closed state** | **Open state** |
| --- | --- | --- |
| **Data collection and processing** |  |  |
| Magnification | 105,000 |  |
| Voltage (kV) | 300 |  |
| Electron dose (e<sup>-</sup>/Å<sup>2</sup>) | 65 |  |
| Under-focus range (μm) | 1.1 - 1.9 |  |
| Pixel size (Å) | 0.828 |  |
| Symmetry imposed | C1 |  |
| Initial particle images (no.) | 844,255 |  |
| Final particle images (no.) | 147,415 | 302,403 |
| Map resolution (Å) | 3.17 | 3.23 |
| FSC threshold | 0.143 | 0.143 |
| Map resolution range (Å) | 2.0 – 10.0 | 2.1 – 13.0 |
| **Refinement** |  |  |
| Initial model used (PDB code) | 1SXJ, 3G65 | Closed state of this study |
| Map sharpening B factor (Å<sup>2</sup>) | –71.6 | –62.8 |
| Map to model CC<sub>mask</sub> | 0.75 | 0.69 |
| Model composition |  |  |
| Non-hydrogen atoms | 21,061 | 20,933 |
| Protein and DNA residues | 2,559; 23 | 2,545; 23 |
| Ligands | 9 | 9 |
| R.m.s. deviations |  |  |
| Bond lengths (Å) | 0.002 | 0.002 |
| Bond angles (°) | 0.552 | 0.609 |
| Validation |  |  |
| MolProbity score | 1.80 | 1.80 |
| Clashscore | 7.36 | 8.43 |
| Poor rotamers (%) | 0.09 | 0.04 |
| Ramachandran plot |  |  |
| Favored (%) | 94.19 | 94.99 |
| Allowed (%) | 5.81 | 5.01 |
| Disallowed (%) | 0 | 0 |

**Extended Data Fig 1.**
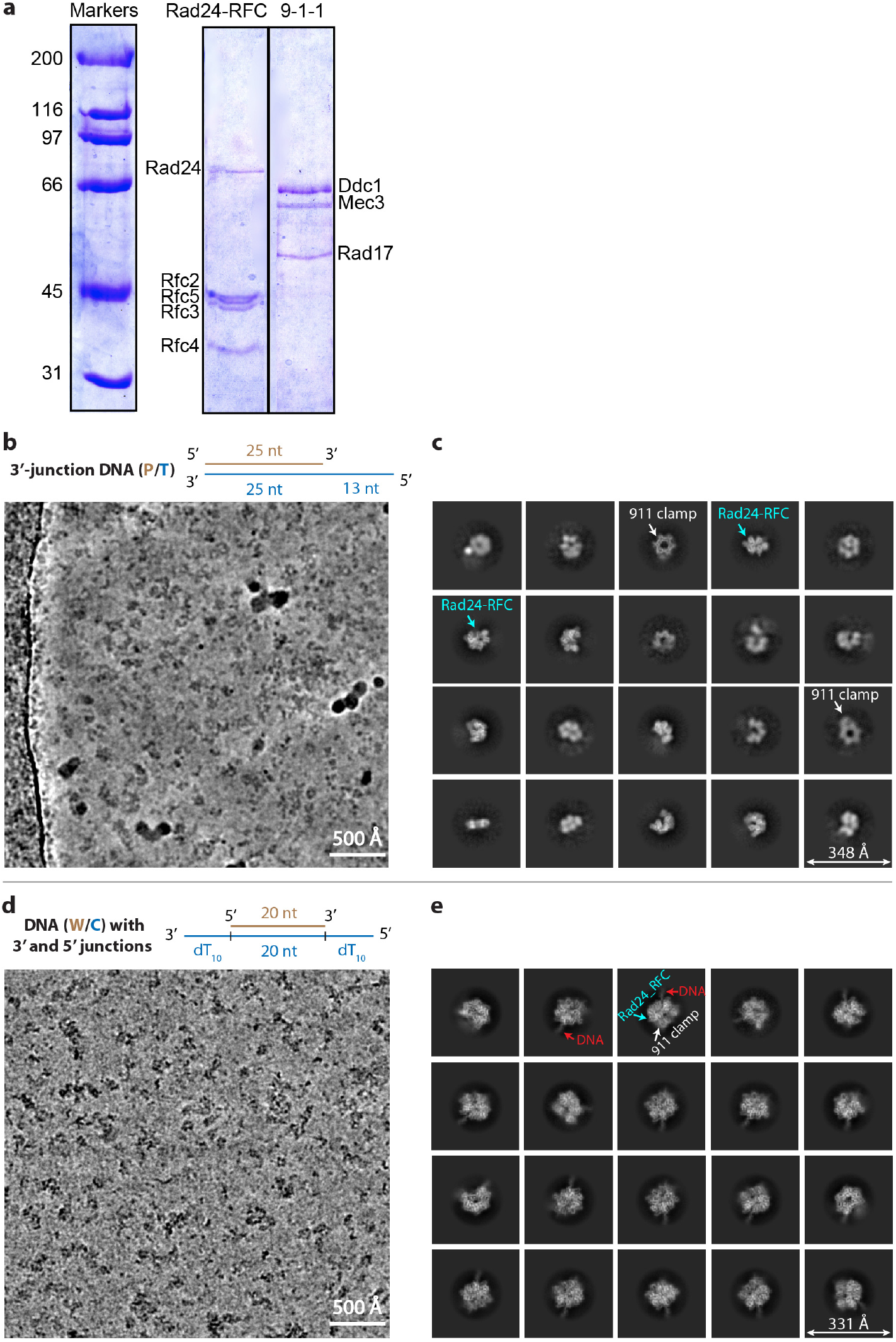
Assembly of the Rad24-RFC–9-1-1–DNA ternary complexes. **a)** SDS-PAGE gels of purified Rad24-RFC and the 9-1-1 clamp. Rad24 is sub stoichiometric. But partial complexes lacking Rad24 were rejected during 2D and 3D image classifications. **b**) A raw micrograph of the mixture of Rad24-RFC with 9-1-1 and a 3’-recessed DNA substrate after incubation in an ice-water bath for 3 hr. **c**) 2D class averages. No ternary complex of Rad24-RFC–9-1-1–DNA was observed. Only the separate 9-1-1 ring and the Rad24-RFC particles were observed. **d**) A typical raw micrograph of the mixture of Rad24-RFC with 9-1-1 and a double tailed DNA substrate containing both the 5’- and 3’-DNA junctions. **e**) Selected 2D class averages with different views, revealing the presence of the targeted Rad24-RFC–9-1-1 clamp–DNA ternary complexes. Scale bars are shown in the lower right corner in each panel.

**Extended Data Fig 2.**
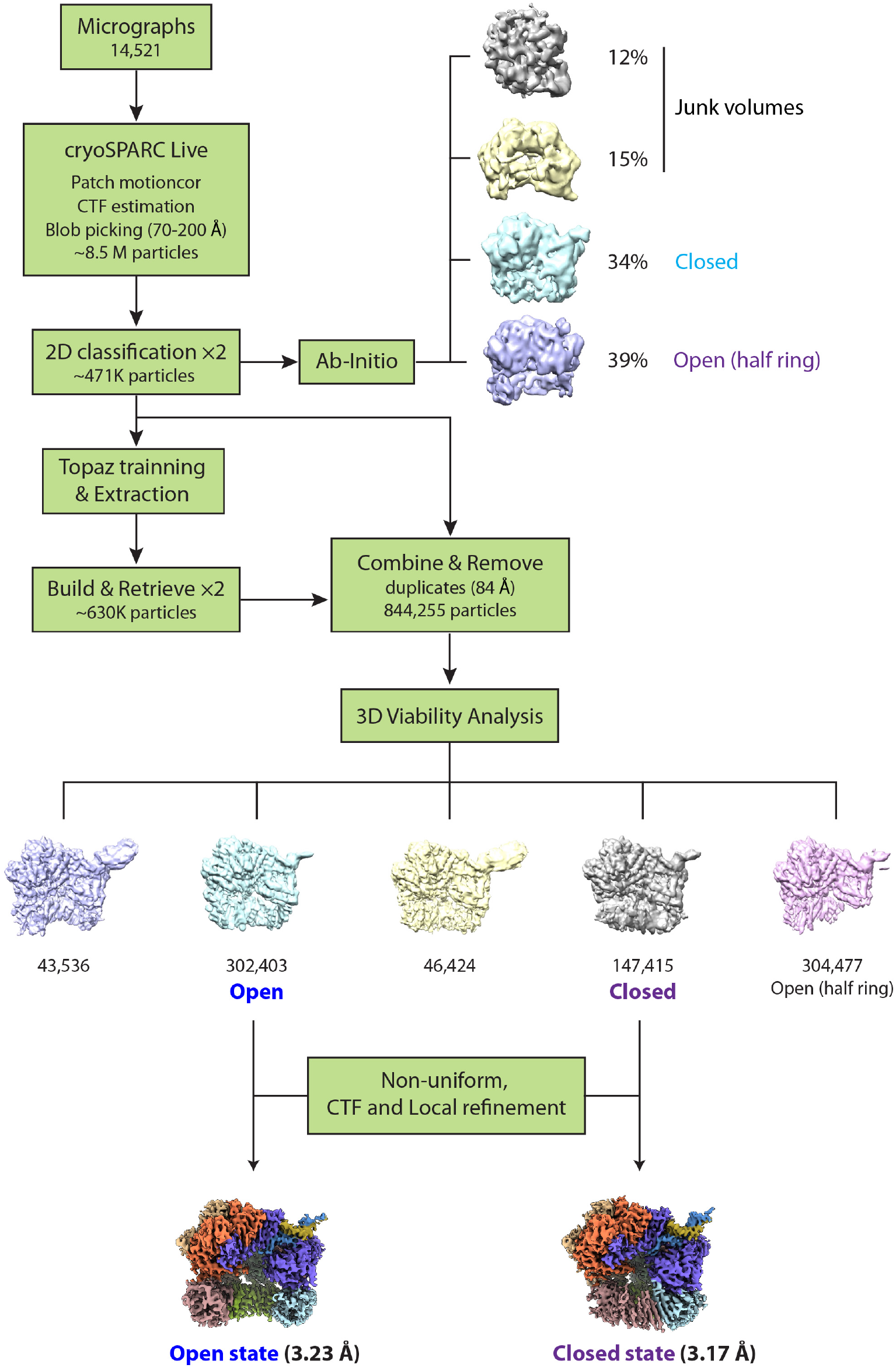
Workflow of image processing and 3D reconstruction. CryoSPARC Live is used to monitor data collection and real-time data processing. We used the particle picking program Topaz (Bepler et al., 2019) and “Build and Retrieve” (Su et al., 2021) to obtain particles with more views. 3D variability analysis was applied to yield the 3.2-Å resolution 3D maps in the closed and open states of the Rad24-RFC–9-1-1 clamp–DNA complex.

**Extended Data Fig 3.**
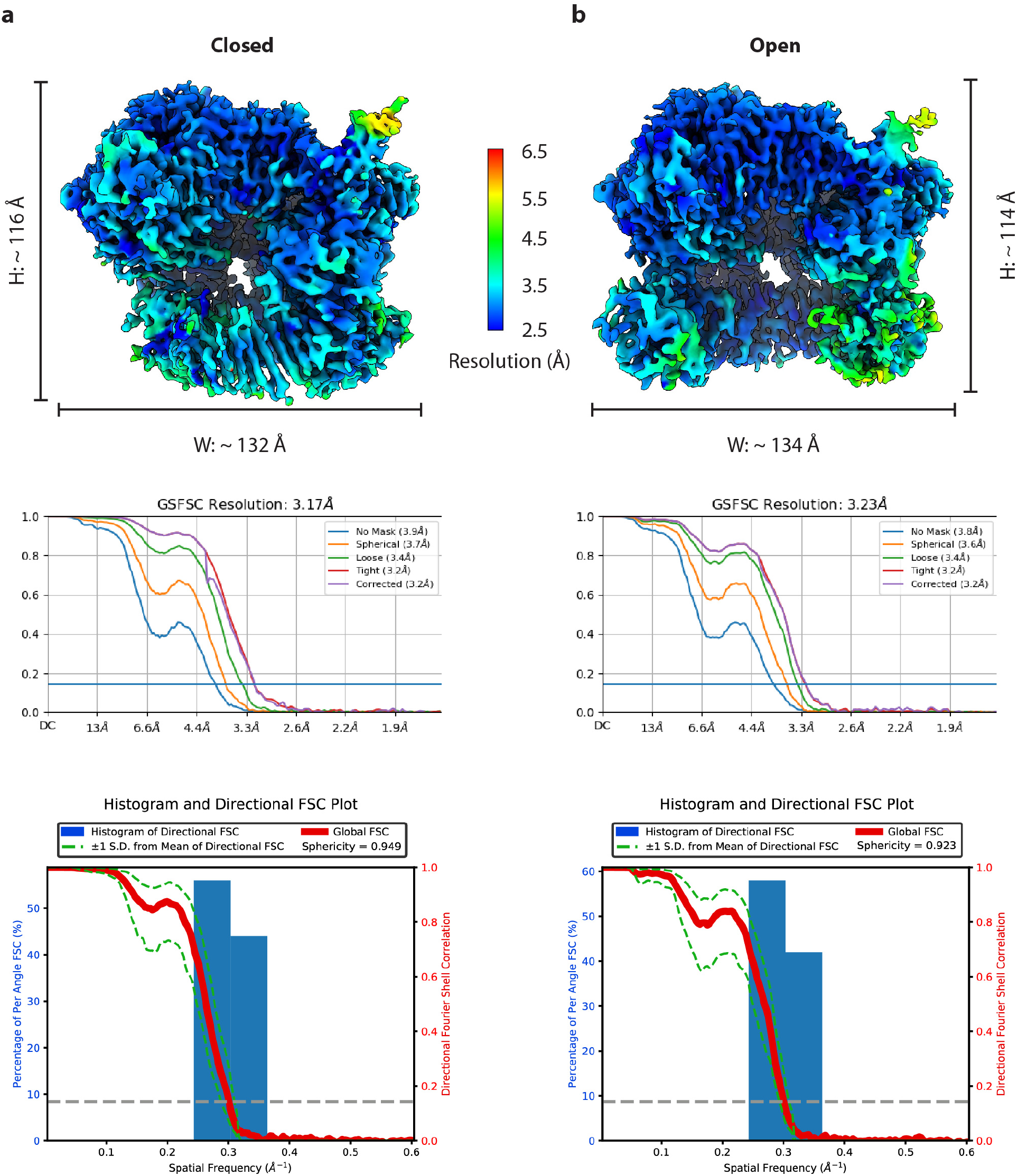
Resolution estimations of the closed (a) and open (b) states of the Rad24-RFC–9-1-1–DNA complex. Upper panels, the two 3D maps colored by local resolution. The dimensions of the structures are labeled. Middle panels, the 0.143 criterion of the gold standard Fourier shell correlation (GSFSC) was used to estimate the average resolutions. Bottom panels, the directional anisotropy of the two maps as quantified by the 3D-FSC server (https://3dfsc.salk.edu/). The sphericity of closed and open state is 0.949 and 0.923, respectively, demonstrating good anisotropic property of both maps.

**Extended Data Fig 4.**
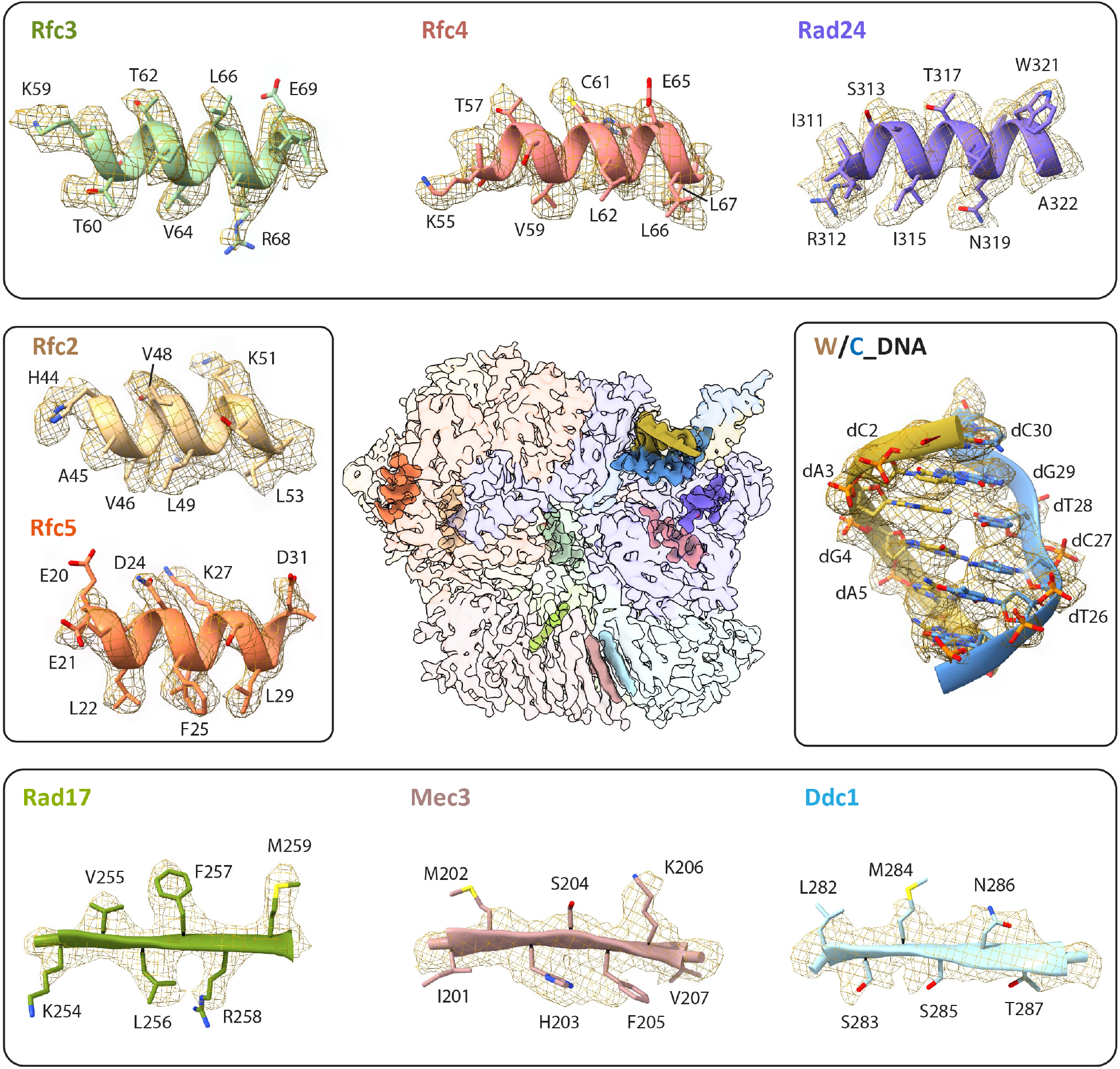
Fitting of the atomic model with the EM density in selected regions of the Rad24-RFC–9-1-1 clamp–DNA complex 3D maps. The central panel shows the segmented 3D map in 90% transparent surface view, except for the nine selected regions shown in solid surface. The nine selected regions – one from each component of the ternary complex – are shown in peripheral panels, with the atomic model in cartoon and residues in sticks, and with the EM density shown in wire meshes. For a fuller representation, the selected upper regions (Rad24-RFC–DNA) used the EM map in the open state, and the selected lower 9-1-1 clamp regions are from the EM map in the closed state.

**Extended Data Fig 5.**
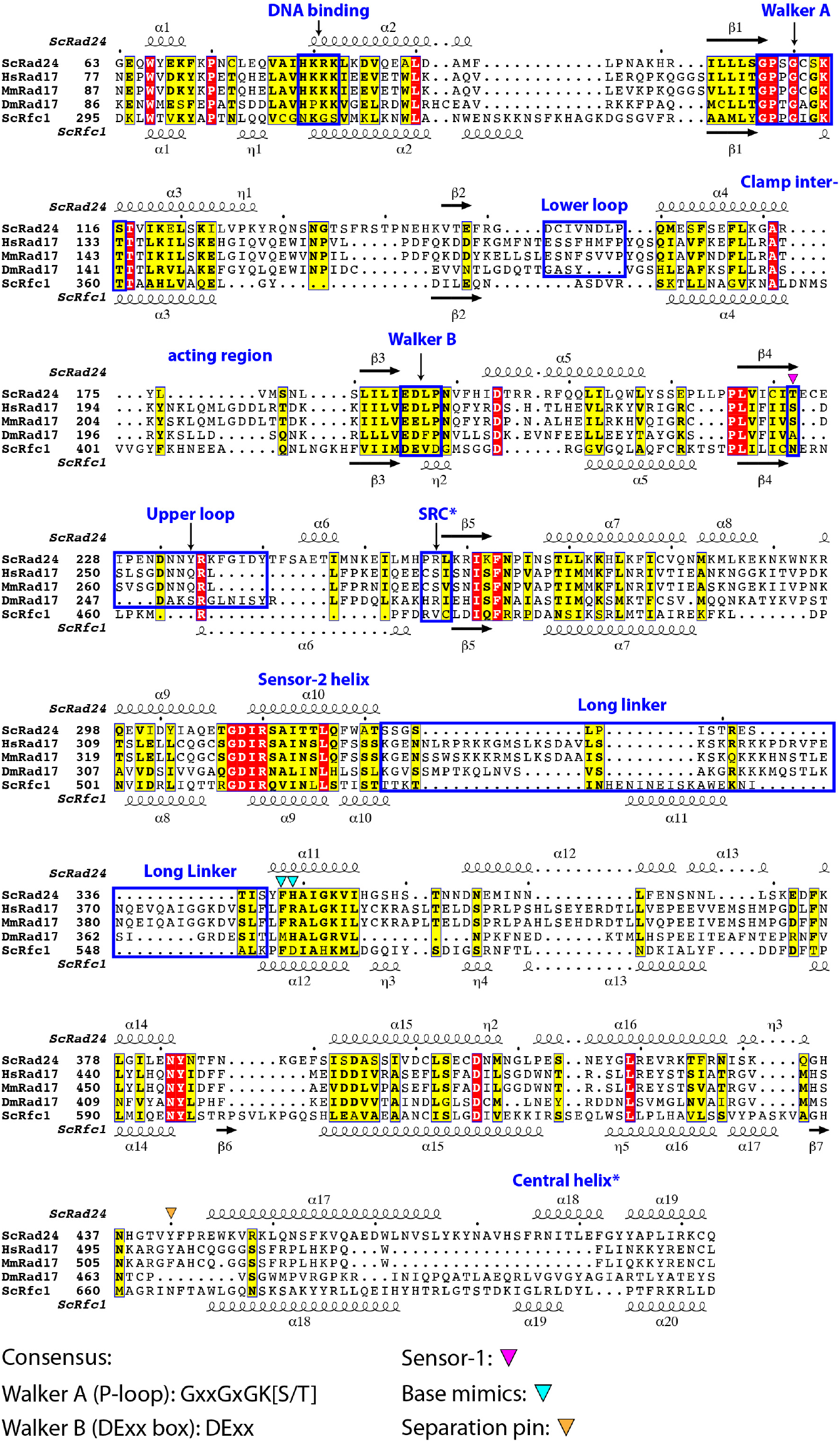
Sequence alignment of yeast Rad24, metazoan Rad17 and yeast Rfc1. Helices and β-strands are shown as coils and arrows, respectively. The assignments are produced by ESPript 3.0 (https://espript.ibcp.fr/ESPript/ESPript/) based on the structures S.c. Rad24 (this study) and S.c. RFC1 (PDB code 1SXJ). The flashy color scheme was applied. Key features are marked above the sequences. The SRC (serine-arginine-cysteine) motif – conserved in Rfc2-5 – is absent in Rad24/RAD17 and Rfc1/RFC1; its location corresponding to the SRC motif in Rfc2-5 is boxed and labeled as SRC*.

**Extended Data Fig. 6.**
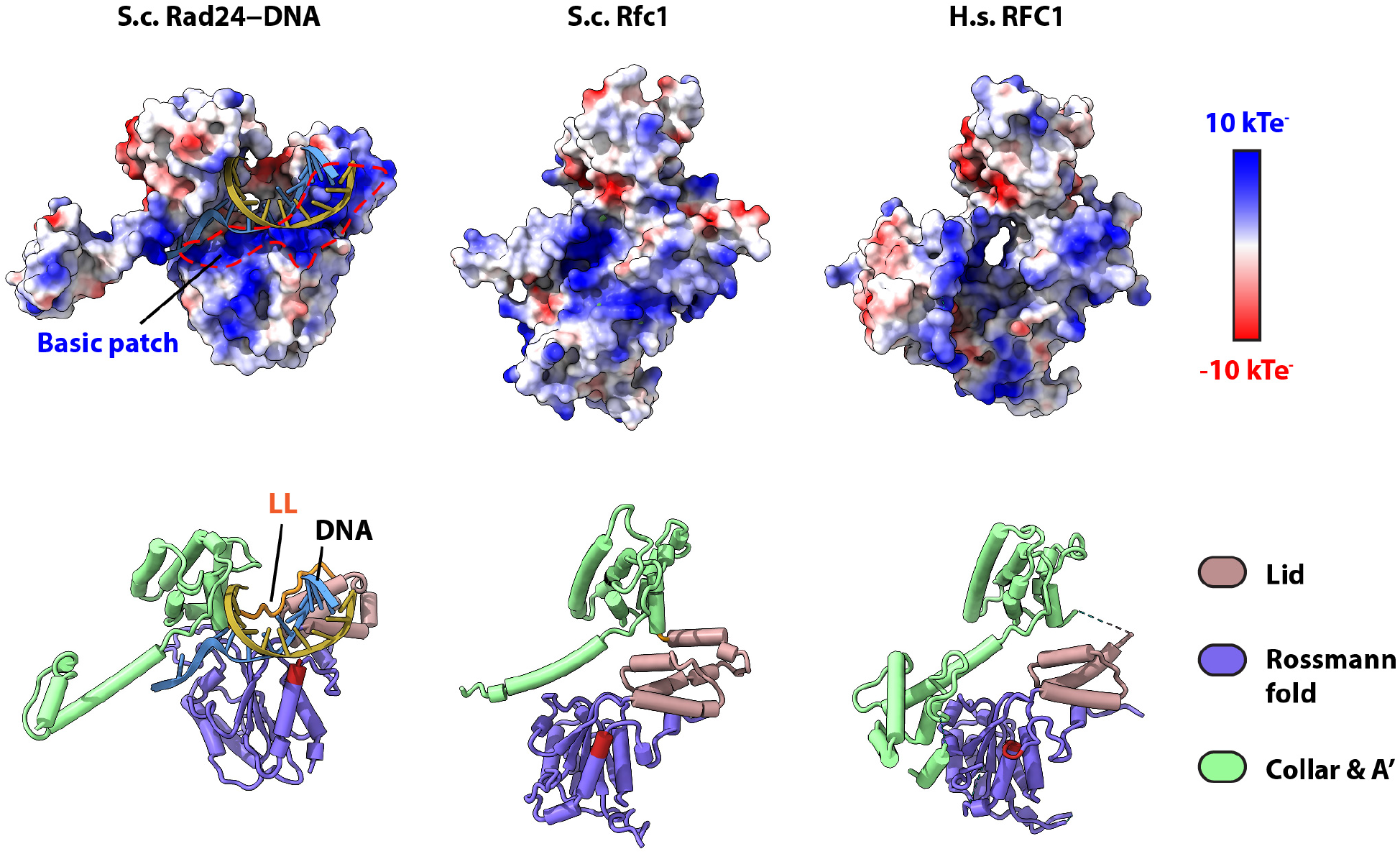
Structural comparison of Rad24 and Rfc1/RFC1. Upper panel, the electrostatic potential charge surface of the S.c. Rad24 (this study), S.c. Rfc1 (PDB code 1SXJ), and human RFC1 (PDB code 6VVO). In Rad24, a contiguous basic patch on top of the AAA+ domain enables DNA binding. Rfc1/RFC1 lack such contiguous basic path. Lower panel, the corresponding atomic models shown in cartoons. These structures are aligned based on the full loader complexes to achieve the best superposition. The four tandem basic residues in Rad24 and the corresponding region in Rfc1/RFC1 are highlighted in firebrick. Note that the lid and Rossmann fold domain move away from the collar domain plus the associated A’ domain by a 35-Å shift and a 20° rotation, creating a large groove for dsDNA binding.

**Extended Data Fig 7.**
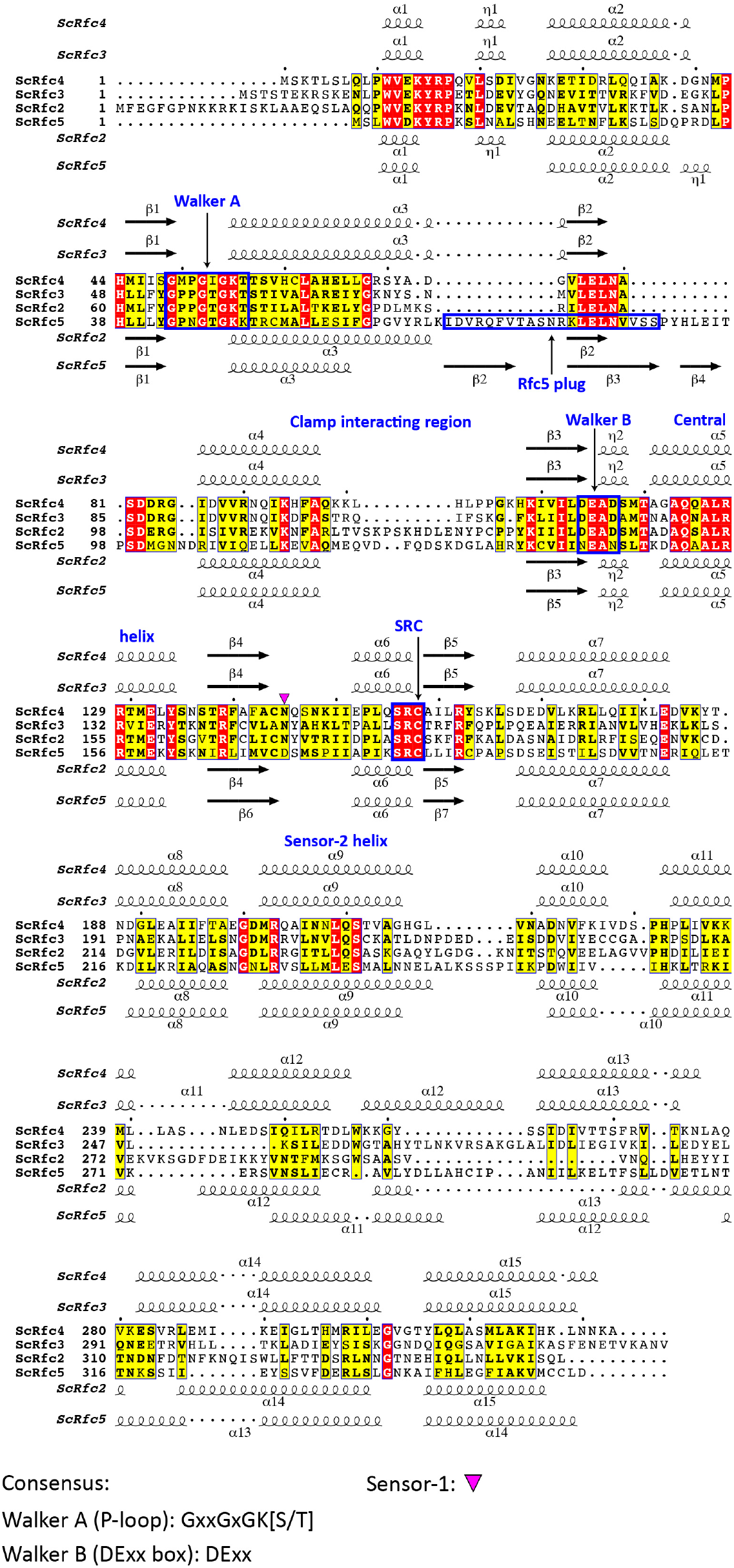
Structure-based sequence alignment of S.c. Rfc2-5. Structural elements of Rfc3 and 4 are shown above and that of Rfc2 and 5 are shown below the sequences. Assignment of the secondary structural elements also considered both the Rad24-RFC structure (current study) and the S.c. RFC structure (PDB code 1SXJ).

**Extended Data Fig 8.**
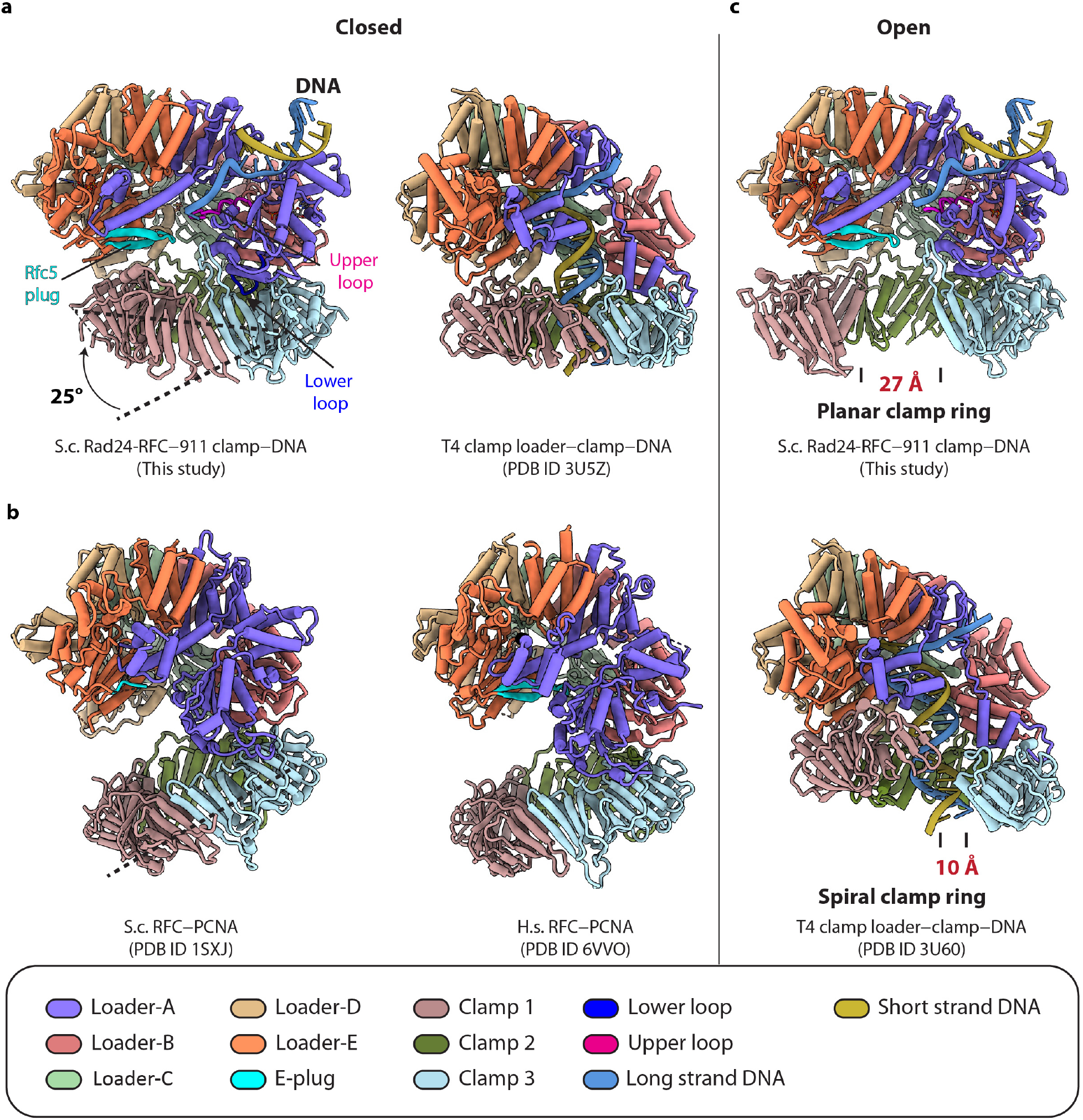
Comparison of the available structures of the DNA clamp–clamp loader–(DNA) complexes. **a**) Structure of the yeast Rad24-RFC bound to 9-1-1- and DNA compared with the T4 phage clamp loader–clamp bound to a DNA substrate. **b**) Comparison of the yeast RFC-PCNA (PDB code 1SXJ) with the human RFC-PCNA (PDB code 6VVO), all in a clamp ring closed configuration. The Rad24-RFC–9-1-1 clamp is more compact than the yeast and human RFC–PCNA structures, with the 9-1-1 ring tilting 25° toward the Rad24-RFC loader. **c**) Structure comparison of the Rad24-RFC–9-1-1 clamp–DNA with the T4 clamp–clamp loader–DNA (PDB code 3U60), both in the open clamp ring conformation. Note the 27-Å DNA gate in the planar 9-1-1 clamp with the 10-Å DNA gate in the T4 spiral clamp. To facilitate comparison, the loader subunits are labeled A through E according to their physical location in the complex.

## LEGENDS OF THE THREE SUPPLEMENTARY VIDEOS

**Supplementary Video 1. Overall structures of the Rad24-RFC–9-1-1 clamp–DNA in the closed and open states.** The closed-state EM map is rotated around the X axis by 360° followed by another 360° rotation around the Y axis. Next, the atomic model is shown morphing between the closed and the open states, highlighting the in-plane rotation of Mec3 that opens a 27-Å gap between Ddc1 and Mec3 of the 9-1-1 clamp. Note that the hook-like lower loop of Rad24 apparently controls the gate opening: the lower loop binds Ddc1 subunit of the 9-1-1 camp in the closed state but becomes flexible and invisible in the open state.

**Supplementary Video 2. Specific interactions between Rad24-RFC and 5’-DNA junction.** DNA is in brown (Watson strand) and steel blue (Crick strand), the Rad24 upper loop in magenta and lower loops in blue, the collar domain a-helix harboring the base mimicking residues in pale green, and the Rfc5 plug in cyan. First, the closed-state atomic model is rotated by 360° around the X axis and then around the Y axis. Next, the scene transits to show the interaction between the tandem basic patch residues 81-HKRK-84 of Rad24 and the dsDNA region. This is followed by a scene showing the Rad24 residues at the 5’-DNA junction. The final scene shows the Rad24 residues that interact and guide the 3’-overhang ssDNA in front of the 5’-junction.

**Supplementary Video 3. Morph between Rfc1 of the S.c. RFC-PCNA structure and Rad24 of the S.c. Rad24-RFC–9-1-1 clamp–DNA structure.** Those two models are aligned by the loaders and shown as cartoons. First, the superimposed models are rotated around the X- and Y-axes. RFC-PCNA is light blue with Rfc1 in salmon. Rad24-RFC–9-1-1 clamp–DNA is in rosy brown with Rad24 in pale green. Next is the morph between the two structures. During morphing, all regions are in transparent gray except for Rfc1 and Rad24. Note that Rad24 shifts 35 Å towards upper right relative to Rfc1 to form a large groove for dsDNA binding. The final scene shows the Rad24 encircling the DNA.

## Notes

### Competing Interest Statement

The authors have declared no competing interest.

